# METTL3 Aggravates Intimal Hyperplasia by Facilitating the m^6^A-YTHDC1-dependent SGK1 Gene Transcription

**DOI:** 10.1101/2025.01.09.632275

**Authors:** Jiaqi Huang, Qianqian Feng, Zhigang Dong, Zhuofan Li, Yihan Liu, Ran Xu, Zhujiang Liu, Qianhui Ding, Xueyuan Yang, Fang Yu, Yiting Jia, Yuan Zhou, Wei Kong, Hao Tang, Yi Fu

**Affiliations:** Department of Physiology and Pathophysiology, School of Basic Medical Sciences, Peking University; State Key Laboratory of Vascular Homeostasis and Remodeling, Peking University, Beijing 100191, China; Department of Neurosurgery, Xuanwu Hospital, Capital Medical University, Beijing 100191, China; China International Neuroscience Institute (China-INI), Beijing, 100191, China; Zhengzhou Key Laboratory of Cardiovascular Aging, National Health Commission Key Laboratory of Cardiovascular Regenerative Medicine, Central-China Fuwai Hospital of Zhengzhou University, Fuwai Central-China Cardiovascular Hospital & Central-China Branch of National Center for Cardiovascular Diseases, Zhengzhou, Henan, 451464, China

**Keywords:** neointimal hyperplasia, vascular smooth muscle cell proliferation, methyltransferase-like 3, serum/glucocorticoid regulated kinase 1, N^6^-methyladenosine

## Abstract

**Background:** Vascular smooth muscle cell (VSMC) migration and proliferation substantially contribute to neointimal hyperplasia related to in-stent restenosis. N^6^-methyladenosine (m^6^A) catalyzed by the methyltransferase-like 3 (METTL3)-involved methyltransferase complex is the most abundant RNA epigenetic modification in eukaryotes, but the role of m^6^A RNA methylation in VSMC migration and proliferation as well as neointima formation remains highly controversial.

**Methods:** Primary human and rat VSMCs were utilized for *in vitro* experiments. VSMC-specific METTL3 knockout mice (*Mettl3*^flox/flox^*Myh11*-CreER^T2^) were generated to explore wire injury in carotid arteries *in vivo*. Methylated RNA immunoprecipitation sequencing (MeRIP-Seq) was performed to screen for target genes by METTL3-catalyzed m^6^A RNA methylation. Methylation site mapping, MeRIP-quantitative PCR (MeRIP-qPCR), chromatin immunoprecipitation-qPCR (ChIP‒qPCR) and reporter gene assays were applied to explore how METTL3 modulates its target gene expression.

**Results:** METTL3 was consistently upregulated in the neointima from mice subjected to carotid wire injury and patients undergoing carotid endarterectomy. VSMC-specific METTL3 deficiency significantly attenuated neointima formation in carotid arteries following wire injury in mice. Accordingly, METTL3 ablation markedly repressed VSMC proliferation both *in vitro* and *in vivo*. Mechanistically, METTL3 directly catalyzed m^6^A methylation on serum/glucocorticoid regulated kinase 1 (SGK1) mRNA and subsequently facilitated its transcription, which relies on the established association between the SGK1 transcript and SGK1 promoter DNA by recruiting the m^6^A reader YTHDC1. Conversely, SGK1 overexpression abolished METTL3 deficiency-mediated suppression of VSMC proliferation and postinjury neointima formation.

**Conclusions:** METTL3-catalyzed m^6^A RNA methylation promoted VSMC proliferation and aggravated postinjury neointima formation by facilitating YTHDC1-dependent SGK1 gene transcription. Targeting the METTL3-YTHDC1-SGK1 axis to modulate VSMC proliferation may be a potential strategy for in-stent restenosis therapy.

## Introduction

Neointimal hyperplasia is a common pathologic feature of atherosclerosis as well as arterial restenosis after angioplasty surgeries such as coronary artery bypass grafting (CABG) and percutaneous coronary intervention (PCI) ^1^. Vascular smooth muscle cell migration and proliferation in response to a myriad of vessel injury-derived stimuli, including biomechanical factors such as stretching, oxidative metabolites, growth factors and cytokines, substantially contributes to neointima formation ^2^. Thus, exploring the underlying regulatory mechanisms of VSMC proliferation and migration could aid in the development of novel therapeutic strategies for treating related vascular pathologies, such as in-stent restenosis and atherosclerosis.

N^6^-methyladenosine (m^6^A) is the most prevalent internal modification of eukaryotic mRNA. This modification is produced by m^6^A “writers”, the major one of which comprises a methyltransferase complex with methyltransferase-like 3 (METTL3), METTL14 and Wilms tumor 1-associated protein (WTAP), as well as additional partner proteins to add m^6^A on RNAs ^3^. METTL3 is the main catalytic subunit with the assistance of METTL14, which stabilizes METTL3 and recognizes target RNAs, while WTAP as a regulatory subunit interacts with the METTL3-METTL14 heterodimer complex and ensures it to be localized in the nuclear spots. Moreover, m^6^A modifications are removed by “erasers”, including fat mass obesity-associated protein (FTO) and alkB homolog 5 RNA demethylase (ALKBH5) and are interpreted by “readers”, such as YTH domain-containing proteins (YTHDF1 – 3;

YTHDC1–2) ^4^. Accumulating evidence has demonstrated that m^6^A modification post-transcriptionally regulates gene expression through specific reader-dependent effects on mRNA splicing, transport, translation, and degradation. Intriguingly, this modification of RNA was recently found to influence its own transcription in a manner dependent on specific “readers” ^5^ . For example, YTHDC1 recognizes enhancer RNA m^6^A methylation to facilitate transcriptional condensate formation and gene transcription, in a scaffold model mediated by YTHDC1 that links the nascent m^6^A-decorated transcript and its promoter DNA via the recognition of m^6^A modifications and its association with transcriptional activators, respectively ^6^. This provides a novel insight into the roles of m^6^A RNA modification.

Current understanding of the regulatory mechanisms required for VSMC migration and proliferation is mainly limited to cell‒cell interactions (e.g., Notch signaling), metalloprotease-mediated extracellular matrix remodeling (e.g., MMP-8 and MMP-9), transcription factor-determined cell identity transition (e.g., NF-kB and KLF4) and microRNA-involved epigenetic modulation ^7–9^. As the most abundant RNA epigenetic modification, the effect of m^6^A RNA methylation on VSMC migration and proliferation as well as neointima formation remains obscure, given that a few studies ^10, 11^, which did not use genetically modified mouse models to verify their findings, present conflicting results.

In this study, we generated VSMC-specific METTL3 knockout mice and found that METTL3 deficiency in VSMCs suppressed postinjury neointima formation *in vivo.* We confirmed that METTL3-catalyzed m^6^A RNA methylation promoted PDGF-BB-induced VSMC proliferation rather than migration. Mechanistically, the recently proposed direct regulation of gene transcription by RNA m^6^A modification was demonstrated to be involved in the modulation of VSMC proliferation; that is, METTL3-mediated m^6^A methylation of SGK1 mRNA facilitates YTHDC1-dependent SGK1 transcription initiation. Targeting the METTL3-YTHDC1-SGK1 axis to suppress VSMC proliferation may be effective in treating restenosis after angioplasty and stenting.

## Materials and methods

### Materials

All Primary antibodies were listed in Supplementary Table I. IRDye-conjugated secondary antibodies for Western blotting analysis were purchased from Rockland, Inc. (Gilbertsville, USA). Secondary antibodies, including 488 anti-mouse (35502) and 633 anti-rabbit (35562) antibodies, for confocal fluorescence microscopy were purchased from Thermo Fisher Scientific (Rochester, USA). Lipofectamine® 3000 Transfection Reagent (L3000015) and Lipofectamine® RNAiMAX Reagent (13778075), which were used for siRNA and plasmid transfection, respectively, were purchased from Invitrogen (Carlsbad, USA). CALNP^TM^ RNAi used for siRNA transfection was purchased from Beijing D-Nano Therapeutics Co., Ltd. (Beijing, China). BrdU (5-bromo-2-deoxyuridine) (19-160) was purchased from Sigma‒Aldrich (St. Louis, MO, USA). 4,6-Diamidino-2-phenylindole (DAPI), the Cell Counting Kit-8 kit (C0039), the BCA reagent (P0011) and the BeyoClick™ EdU-488 proliferation kit (C0071) were purchased from Beyotime Biotechnology (Shanghai, China). Smooth muscle cell medium (1101) was obtained from ScienCell (Carlsbad, CA, USA). Dulbecco’s modified Eagle’s medium (DMEM) and Fetal Bovine Serum (FBS) were obtained from Gibco. (NY, USA).

### Human Samples

The human samples of plaque and control artery were obtained from China Carotid Atherosclerosis Biobank Consortium (CCABC), Xuanwu Hospital, Capital Medical University (Beijing, China). This study involving human tissue was approved by the Medial Ethics Committee of Xuanwu Hospital (No. 2021124). Normal carotid arteries were obtained from people who died accidentally as control samples. Patient carotid arteries were collected from patients undergoing human carotid endarterectomy. The clinical characteristics of the patients are shown in Supplementary Table II. The samples of both normal carotid arteries and patient plaques were used for protein or mRNA isolation as well as immunofluorescent staining.

### Mouse Carotid Artery Wire Injury

Carotid artery wire injury surgery was performed on 8-week-old male mice ^12^. Briefly, after isolation of the left common carotid artery, 6-0 silk slipknots were used to block the blood flow of the common carotid artery and the internal carotid artery. Next, a curved flexible wire (0.014 inches, approximately 0.35-0.38 mm) with 1% heparin was inserted into the external carotid artery to induce carotid artery injury. Then, the wire was removed, and the left external carotid artery was tied proximally with 6-0 silk. The skin incision was closed. 7 days, 14 days or 28 days after injury, the mice were sacrificed, and the right carotid artery was isolated as uninjured control. After removal of the perivascular tissues, the carotid arteries were harvested for further Western blotting analysis. Some of the carotid arteries were collected for H&E staining and neointima area analysis.

### Western Blotting Analysis

Western blotting analysis was performed as previously described ^13^. Briefly, cell lysates and tissue extract of equal total proteins quantified by BCA reagent were resolved by SDS‒PAGE followed by transfer to nitrocellulose membranes. Then, the membranes were blocked using 5% BSA or milk in 1× TBST for 2 hours. After incubation with primary antibodies at 4 °C overnight on a horizontal shaker, the membranes were washed with 1× TBST 3 times. The membranes were treated with IRDye-conjugated secondary antibodies (Rockland, Inc., Gilbertsville, PA, USA) for 1 hour at room temperature. After the nonspecific antibody was washed out with 1× TBST 3 times, the immunofluorescence signals were detected by using an Odyssey infrared imaging system (LI-COR Biosciences, Lincoln, NE, USA).

### RT‒qPCR Analysis

RT‒qPCR analysis was performed by using an ABI QuantStudioTM 3 Real-Time PCR System (Rockford, CA, USA). The 18S ribosomal RNA level was used as the internal control for normalization. For human atherosclerotic carotid arteries, total RNA was extracted from control carotid arteries and neointimal tissues from patients undergoing human carotid endarterectomy (CEA). For analysis of the expression of METTL3, SGK1 or pre-SGK1, total RNA was extracted from cultured human and rat VSMCs. cDNA was obtained by using cDNA Synthesis SuperMix (E047-01B, Novoprotein) (Suzhou, China) according to the manufacturer’s instructions. Then, RT‒ qPCR analysis was performed by using specific primers. The primers used for all RT‒ qPCR analyses are presented in Supplementary Table III.

### Mice

*Mettl3^flox/flox^* mice were kindly provided by Prof. Wengong Wang from Peking University. *Myh11-*CreER^T2^ mice were obtained from the Jackson Laboratory (Strain: 019079). For Cre-inducible mice, *Myh11-*CreER^T2^ mice were crossed with *Mettl3^flox/flox^*mice. Since *Myh11-*CreER^T2^ mice were created by the insertion of the bacterial artificial chromosome containing *Myh11*-CreER^T2^ in the Y chromosome, *Myh11-*CreER^T2^ genotypes were limited in male mice. In accordance, only male mice were applied in current study^14^. Eight-week-old male *Mettl3^flox/flox^*; *Myh11-*CreER^T2^ mice were treated with 75 mg/kg tamoxifen for 5 consecutive days to induce VSMC-specific *Mettl3* knockout. *Myh11*-CreER^T2^ mice with tamoxifen injection, *Mettl3*^flox/flox^ with Tamoxifen injection, and *Mettl3*^flox/flox^*; Myh11*-CreER^T2^ with corn oil injection mice were used as controls. The primers used for genotyping were listed in Supplementary Table IV. Both *Mettl3^flox/flox^*and *Myh11-*CreER^T2^ mice were maintained on a C57BL/6 strain background. All mouse experiments were approved (LA2021125) by the Institutional Animal Care and Use Committee of Peking University Health Science Center, and the care of the animals was in accordance with institutional guidelines.

### Measurement of Blood Pressure

Blood pressure levels were recorded by using a CODA Mouse & Rat Tail-Cuff Blood Pressure System (Kent Scientific Co., Connecticut, USA). Briefly, mice or rats were placed in the restraint corridor and allowed at least 10 minutes of acclimation. The area was warmed with a heating pad and a quiet, dark environment was maintained to ensure reliable measurements within the parameters of this technology. The blood pressure of each mouse or rat was tested for 6 consecutive times to calculate the mean value.

### Morphometric Analysis and Quantification of Neointima Formation

Mice were euthanized by CO_2_ inhalation, perfused with PBS and fixed with 4% paraformaldehyde. Arterial specimens were subjected to OCT embedding and sectioning. Serial sections (6 μm) were collected at 100 μm intervals (10 sections per segment/interval), mounted on slides, and numbered. Six digitized sections with the same identification number from three segments/intervals (∼0.4 mm, 0.5 mm, and 0.6 mm from the injury site) of each animal were stained with H&E for morphometric analysis. The procedure for neointima quantification was similar to previous descriptions ^15^. Briefly, the areas of the media and intima on cross-section of H&E-stained artery segments were automatically measured in square pixels (pixel^2^) with a computerized image analysis system (Image-Pro Plus 6.0 software) by two experienced investigators blinded to the treatments.

### Cell Culture

Primary rat VSMCs were isolated from the thoracic aortas of male Sprague‒ Dawley rats (weight 150‒180 g) by collagenase digestion as previously described ^12^. Primary VSMCs were cultured in low-glucose Dulbecco’s modified Eagle’s medium (DMEM) (Gibco BRL, Grand Island, NY, USA) supplemented with 10-20% FBS (Gibco BRL, Grand Island, NY, USA), and cells before passage 8 were used for further experiments.

The human umbilical cord for isolating human umbilical artery smooth muscle cells were obtained from Peking University People’s Hospital (Beijing, China). The procedures for the isolation of human umbilical artery SMCs were approved by the Ethics Committee of the Peking University Health Science Center (2015PHB024). The cells were digested and passaged with 0.05% trypsin. Primary human umbilical artery SMCs within passages 4-7 were cultured in smooth muscle cell medium supplemented with 2% FBS, 1% smooth muscle cell growth supplement (SMCGS) and 1% penicillin/streptomycin as previous reported^16^. Human umbilical artery SMCs were verified by SMA immunofluorescence staining (Supplementary Figure 1).

The rat aortic smooth muscle cell line A7r5 (CRL-1444) was cultured in high-glucose DMEM supplemented with 10% FBS, 100 units/mL penicillin and 100 μg/mL streptomycin. All cells were maintained in a humidified 5% CO_2_ incubator at 37 °C.

### Small Interfering RNA Transfection

The siRNAs were designed and synthesized by GenePharma (Suzhou, China). *In vitro* transfection of primary rat and human VSMCs with siRNAs (20 nM) was performed by using RNAiMAX reagent (Invitrogen, CA, USA) according to the manufacturer’s protocol. The sequences of the siRNAs used are listed in Supplementary Table V.

### Measurement of VSMC proliferation

Cell Counting Kit-8 (CCK-8) and EdU assays were used to determine cell proliferation as previously described ^13^. After 48 hours, primary rat or human VSMCs transfected with scramble siRNA or METTL3 or YTHDC1 siRNA were seeded at 5×10^3^ cells/well in a 96-well culture plate with FBS-free medium for 12 hours. Then, the medium was replaced with medium containing 20 ng/mL of PDGF-BB. Twenty-four hours later, CCK-8 solution was added to the cells for 2 hours, and the optical density (OD) of the cells was measured at 450 nm on a microplate reader. Cells without CCK-8 solution were used as a blank control. For the EdU assay, PDGF-BB-treated cells were incubated with EdU working reagent for 2 hours and then fixed with 4% PFA for 15 minutes. EdU incorporation was detected with a BeyoClick™ EdU-488 Kit, and images were captured with a BioTek Cytation 5 (Agilent, VT, USA). Image-Pro Plus 6.0 software was used to count the number of cells. For cell cycle assay, the Cell Cycle Analysis Kit (C1052; Beyotime, Shanghai, China) was used. Rat VSMCs were collected and then stained with a solution containing propidium iodide (0.05 mg/mL), RNase A (1 mg/mL), and 0.3% Triton X-100 in the dark for 30 min. The percentage of VSMCs in different cell cycle phases was examined by measuring the DNA content (propidium iodide intensity) with a flow cytometer (Beckman Coulter, Brea, CA), and populations of G1, S, and G2/M phase cells were determined with the ModFIT software.

### Measurement of VSMC Migration

Transwell migration assays and wound healing assays were used to determine cell migration as previously described ^15^. After 48 hours, 5×10^4^ primary rat or human VSMCs transfected with scramble siRNA or METTL3 siRNA were seeded in the top chambers of Transwell plates (BD Biosciences, San Diego, CA, USA) in FBS-free DMEM with a 8 µm pore size membrane inserted. The lower compartment contained DMEM supplemented with 20 ng/mL of PDGF-BB. Twelve hours later, the cells that migrated through the membrane to the bottom chamber were fixed with 4% PFA for 15 minutes. The cells were washed twice with PBS. Then, the cells in the upper compartment of the insert were gently removed by wiping the upper side of the membrane with a cotton swab. The cells on the lower surface of the membrane were stained with 0.1% crystal violet staining solution for 15 minutes, after which the cells were washed twice with PBS. Then, images were captured by microscopy. Next, the crystal violet was completely eluted with 33% acetic acid, and the OD of the eluate was measured at 570 nm and 630 nm on a microplate reader, which indirectly reflects the number of cells. For the wound healing assay, primary rat VSMCs transfected with scrambled siRNA or METTL3 siRNA were seeded in a 6-well plate after 48 hours and wounded by manually scraping the cells using a 200 µL pipette tip. The cells were cultured in FBS-free DMEM for 12 hours, and the medium was then replaced with DMEM containing 20 ng/mL of PDFG-BB. The migration areas were monitored by microscopy. Image-Pro Plus 6.0 software was used to measure the migration area.

### Measurement of VSMC apoptosis

For cell apoptosis detection, the crossed-sections or VSMCs were detected by terminal deoxynucleotidyl transferase-mediated dUTP-biotin nick end labeling (TUNEL) assay (C1086, Beyotime, China) according to the manufacturer’s instructions.

### BrdU Incorporation Assay

A carotid artery wire injury mouse model was established in 8-week-old male *Mettl3* ^SMCWT^ mice and *Mettl3*^SMCKO^ mice as described above ^12^. Fourteen days after the surgery, the mice were sacrificed. The mice were intraperitoneally injected with two subcutaneous doses of BrdU (100 mg/kg) at 18 hours and 3 hours before they were sacrificed. Then, the artery was embedded in OCT compound after fixation with 4% PFA. Artery sections were washed with PBS and incubated with 0.1% Triton-X 100 for 20 minutes. The samples were neutralized by incubation in phosphate/citric acid buffer (pH 7.4) for 20 minutes and then blocked with 3% BSA for 1 hour. Anti-BrdU antibody (66241-1-Ig) was incubated at a 1:200 dilution overnight. After washing with PBS, the sections were incubated with secondary antibodies for 1 hour. The nuclei were stained with DAPI.

### Immunofluorescence Staining

Immunofluorescence staining was performed as previously described ^12^. Briefly, isolated mouse carotid artery samples were fixed with 4% PFA overnight and then embedded in OCT compound. Then, 4-8-µm-thick sections were generated using a cryostat. Before staining, the sections were treated at room temperature for 30 minutes. Following the treatment with 0.1% Tween 20-PBS for 5 minutes, the sections were incubated with normal IgG antibodies or an anti-SGK1 antibody (Santa Cruz) at 1:200 or anti-smooth muscle actin (SMA) antibody at 1:400 in PBS overnight at 4 °C. After 3 washes with PBS, the sections were then incubated with secondary antibody at a 1:1000 dilution in PBS at room temperature for 1 hour. Then, the sections were incubated with DAPI at a 1:1000 dilution for 5 minutes. For human samples, we used multiplex Immunohistochemistry (mIHC) staining with the AlphaTSA Multiplex IHC Kit (AXT6110000, AXT6410000, AXT6610000, Alpha X Biotech, China) by AlphaXPainter X30. Briefly, All the primary antibodies were incubated for 1 hour at 37°C. Then slides were incubated with Alpha X Ploymer HRP Ms+Rb (Alpha X Bio, Beijing, China) for 10 minutes at 37°C. Alpha X 4-Color IHC Kit (AXT37100031, Alpha X Bio, Beijing, China) was used for visualization. After each staining cycle, heat-induced epitope retrieval was performed to remove all the antibodies including primary & secondary antibodies. The slides were counter-stained with DAPI for 5 minutes and enclosed in Antifade Mounting Medium (I0052; NobleRyder, Beijing, China). The immunofluorescence signals were detected using a Leica TCS SP8 CARS system and FV3000. Images were analyzed with Image J software.

### MeRIP Sequencing

MeRIP experiments, high-throughput sequencing and data analysis were conducted by SeqHealth Technology Co., Ltd. (Wuhan, China). Briefly, total RNA was extracted from control or METTL3-overexpressing rat primary VSMCs using TRIzol Reagent (Invitrogen, cat. No. 15596026). One hundred micrograms of total RNA were used for the MeRIP experiment. Briefly, 20 mM of ZnCl_2_ was added to the total RNA and incubated at 94 °C for 5 minutes until the RNA fragments were mainly 100-200 nt in length. Then, 10% of the RNA fragments were saved as “input”, and the remaining fragments were used for m^6^A immunoprecipitation (IP). A specific anti-m^6^A antibody (Synaptic Systems, 202003) was used for m^6^A IP. The stranded RNA sequencing library was constructed by using the KC-Digital^TM^ Stranded mRNA Library Prep Kit for Illumina® (Catalog No. DR08502, Wuhan Seqhealth Co., Ltd., China) with five PCR cycles. The kit eliminates duplication bias in the PCR and sequencing steps. Then, ribosomal cDNA was removed by a SMARTer Stranded Total RNA-Seq Kit version 2 (Pico Input Mammalian; 634413; TaKaRa/Clontech, Japan), and ten PCR cycles were added. The library products corresponding to the 200-500 bp fragments were enriched, quantified and finally sequenced on a DNBSEQ-T7 sequencer (MGI Tech Co., Ltd., China) with a PE150 model. The datasets produced in this study are available in the following databases: Gene Expression Omnibus GSE252691 (https://www.ncbi.nlm.nih.gov/geo/query/acc.cgi?&acc=GSE252691).

### MeRIP-qPCR

m^6^A modifications of individual genes were determined using the MeRIP‒qPCR assays according to the SYSY m^6^A antibody manufacturer’s instructions. Briefly, total RNA was extracted from control or METTL3 knockdown rat primary VSMCs using TRIzol Reagent (Invitrogen, cat. no. 15596026), and 100 μg of total RNA was used for the MeRIP‒qPCR experiment. ZnCl_2_ (20 mM) was added to the total RNA and incubated at 90 °C for 5 minutes until the RNA fragments were mainly distributed at 100–200 nt. Then, one tenth of the RNA fragments were saved as “input”, and the remaining fragments were used for m^6^A IP. The RNA was incubated with 5 μg of anti-m^6^A antibody (202,003, Synaptic Systems) or rabbit IgG overnight at 4 °C with rotation with RNase inhibitors in 1× immunoprecipitation buffer. Then, 60 µL of Pierce™ Protein A/G Magnetic Beads (88,803, Thermo Scientific) was prewashed and mixed with each antibody-conjugated RNA tube for 4 hours at 4 °C with rotation. After 3-5 washes, the methylated mRNAs were precipitated with 20 µg of GlycoBlue™ and one-tenth volumes of 3 M sodium acetate in 2.5 volumes of 100% ethanol at −80 °C overnight. Further enrichment was calculated by qPCR, and the corresponding m^6^A enrichment in each sample was calculated by normalization to the input. The primers used for MeRIP‒qPCR analyses for SGK1 are presented in Supplementary Table VI.

### RNA Stability Assay

Primary rat VSMCs were transfected with siRNA using RNAiMAX reagent (Invitrogen) and then seeded on 6-well plates. For inhibition of transcription, cells were cultured in medium supplemented with 2 µg/mL of actinomycin D and harvested at 0, 2, 4, 6, and 8 hours after treatment. RNA was extracted with TRIzol and subjected to RT-qPCR analysis. The half-life was calculated as previous reported^17, 18^. Briefly, we first analyzed the qPCR data and normalized the target Ct average of each time point to the internal reference (GAPDH) Ct average to obtain the ΔCt value (ΔCt = (target Ct of each time point − internal reference Ct of the same time point)). Then we normalized the ΔCt value of each time point to the ΔCt value of t=0 to obtain the ΔΔCt value (ΔΔCt = (ΔCt of each time point − ΔCt of t=0)). We then calculated the relative RNA quantity for each time point as RNA quantity = 2^(−ΔΔCT). To further analyze the data, we determined the RNA decay rate through non-linear regression curve fitting, specifically employing a one-phase decay model in GraphPad Prism. In detail, the fitting process utilized the least squares method with an ordinary fit, maintaining a confidence level of 95%. Additionally, we opted for asymmetrical confidence intervals (CIs) to enhance the robustness of our results. The goodness of fit was quantified using the R-squared value, while convergence criteria were set to medium to ensure reliable fitting. The half-life (t_1/2_) of SGK1 mRNA was fitted and calculated independently for each experiment and statistical analyzed by unpaired two-tailed Student’s *t* test.

### RNA Immunoprecipitation (RIP)

A7r5 cells were transfected with Flag-tagged YTHDC1 using Lipofectamine 3000 reagent (Invitrogen), divided into two equal groups, and then transfected with siRNAs using RNAiMAX reagent (Invitrogen). After UV crosslinking, the cells were harvested with a cell scraper and lysed with RIP lysis buffer (50 mM HEPES, 150 mM KCl, 2 mM EDTA, 1% NP40, pH 7.5, protease inhibitor, and 1 U/µL RNase inhibitor). The cell lysates were incubated with 3 µg of Flag antibody (Millipore Sigma) or normal antibodies against mouse IgG (Macgene) overnight at 4 °C and then incubated with magnetic beads (Thermo Fisher Scientific) for 5 hours at 4 °C with orbital rotation. Immunoprecipitated samples were adequately washed with RIP-wash buffer (50 mM HEPES, 300 mM KCl, 0.05% NP40, pH 7.5) and subjected to DNase I (Thermo Fisher Scientific) digestion and then protease K (Tiangen, supplemented with 1% SDS) digestion in NT2 buffer (50 mM Tris-HCl, 150 mM NaCl, 1 mM MgCl_2_, 0.05% NP40, 1 U/µL RNase inhibitor, pH 7.5). RNA was extracted with TRIzol and subjected to RT‒ qPCR analysis. The enrichment of immunoprecipitated RNA was normalized to the input.

### Chromatin Immunoprecipitation (ChIP)

A7r5 cells were transfected with Flag-tagged YTHDC1 using Lipofectamine 3000 (Invitrogen), divided into two equal groups, and then transfected with siRNAs using RNAiMAX (Invitrogen). Cells were crosslinked with 1% formaldehyde, which was stopped by the addition of 0.125 M glycine. Cells were harvested with a cell scraper and lysed with ChIP lysis buffer (0.5% SDS, 10 mM EDTA, 50 mM Tris-HCl, pH 8.0, protease inhibitor). The cell lysate was sonicated to fragment the chromosomal DNA. After centrifugation, the supernatant was precleared with magnetic beads (Thermo Fisher Scientific) for 2 hours at 4 °C using orbital rotation. Moreover, magnetic beads were incubated with 3 µg of Flag antibody or normal antibodies against mouse IgG (Macgene) in ChIP IP buffer (20 mM HEPES, 200 mM NaCl, 2 mM EDTA, 0.1% Na-DOC, 1% Triton X-100, 5 mg/mL BSA, pH 8.0, protease inhibitor) at room temperature for 2 hours, after which a quarter volume of ChIP lysate was added for immunoprecipitation overnight at 4 °C. Immunoprecipitated samples were washed 4 times in LiCl wash buffer (50 mM HEPES, 1 mM EDTA, 1% NP40, 1% Na-DOC, 500 mM LiCl, pH 8.0) and once in TE buffer (10 mM Tris HCl, 1 mM EDTA, pH 8.0). Immunoprecipitated samples were eluted twice with elution buffer (50 mM NaHCO_3_, 1% SDS) at 37 °C and shaken at 1000 rpm for 30 minutes. The samples were dispersed in elution buffer with 0.45 M NaCl at 65 °C, shaken at 1200 rpm for 2 hours and digested with proteinase K at 65 °C for 15 minutes. DNA was isolated by the phenol‒ chloroform extraction method and determined by regular PCR or qPCR analysis. The enrichment of immunoprecipitated DNA was normalized to the input. The primers used for ChIP‒PCR analyses for SGK1 promoter fragment are presented in Supplementary Table VII.

### Dual-luciferase Reporter Assay

A7r5 cells were transfected with siRNA using RNAiMAX reagent (Invitrogen), seeded on 6-well plates and cotransfected with wild-type or mutated pGL3-basic reporter plasmid (SGK1-wt, SGK1-Ddel, SGK1-Rmut or SGK1-DdelRmut) and Renilla reporter plasmid using PEI reagent. After 48 hours, firefly and Renilla luciferase activities were measured using a Dual Luciferase Reporter Assay Kit (Vazyme, Nanjing, China) according to the manufacturer’s instructions. Relative fold changes of luminescence units (RLUs) catalyzed by firefly luciferase to RLUs catalyzed by Renilla luciferase were calculated.

### Adeno-associated virus infection

To specifically overexpress SGK1 in VSMCs in mice, recombinant adeno-virus serotype 9 (AAV-9) vectors with the SM22α promoter carrying GFP controls or full-length SGK1 were manufactured by Genechem Co., Ltd. (Shanghai, China). Then, 10-week-old male *Mettl3* ^SMCWT^ mice and *Mettl3*^SMCKO^ mice were injected with either AAV-SM22α-GFP control (2.5×10^12^ vg/mouse) or AAV-SM22α-SGK1 (2.5 ×10^12^ vg/mouse). Two weeks later, carotid artery wire injury was induced as described above.

### Statistical analysis

All the data are presented as the mean ± standard error of the mean (SEM). Statistical analysis was performed using GraphPad Prism 8.0 software (GraphPad Software, San Diego, CA, USA). For normally distributed data, unpaired Student’s *t* test was utilized for two-group comparisons, whereas ANOVA followed by post hoc comparisons was used for multiple group data analyses. In all analyses, a *P* value < 0.05 was regarded as statistically significant. The detailed statistical analyses used for each experiment are presented in the corresponding Figure legends.

## Results

### METTL3 upregulation is correlated with neointimal hyperplasia

We collected neointimal tissues from patients who underwent carotid endarterectomy and detected the expression of m^6^A methyltransferase complex components (METTL3, METTL14, and WTAP) and demethyltransferase (ALKBH5 and FTO). As results, METTL3 was exclusively upregulated in neointima compared with the control arterial tissues (Figure 1A-B). Immunofluorescence staining further confirmed METTL3 protein was mainly upregulated in VSMCs in the human neointima (Figure 1C and Supplementary Figure 2A). In addition, we validated the gradual upregulation of METTL3 in carotid arteries at different stages following wire injury in mice (Figure 1D), while the upregulated METTL3 was mainly expressed in VSMCs (Figure 1E). Since METTL3 is the core catalytic subunit of the methyltransferase complex, these data collectively suggested that the increase of METTL3-catalyzed m^6^A RNA methylation in VSMCs is probably correlated with neointimal hyperplasia.

**Figure 1.**
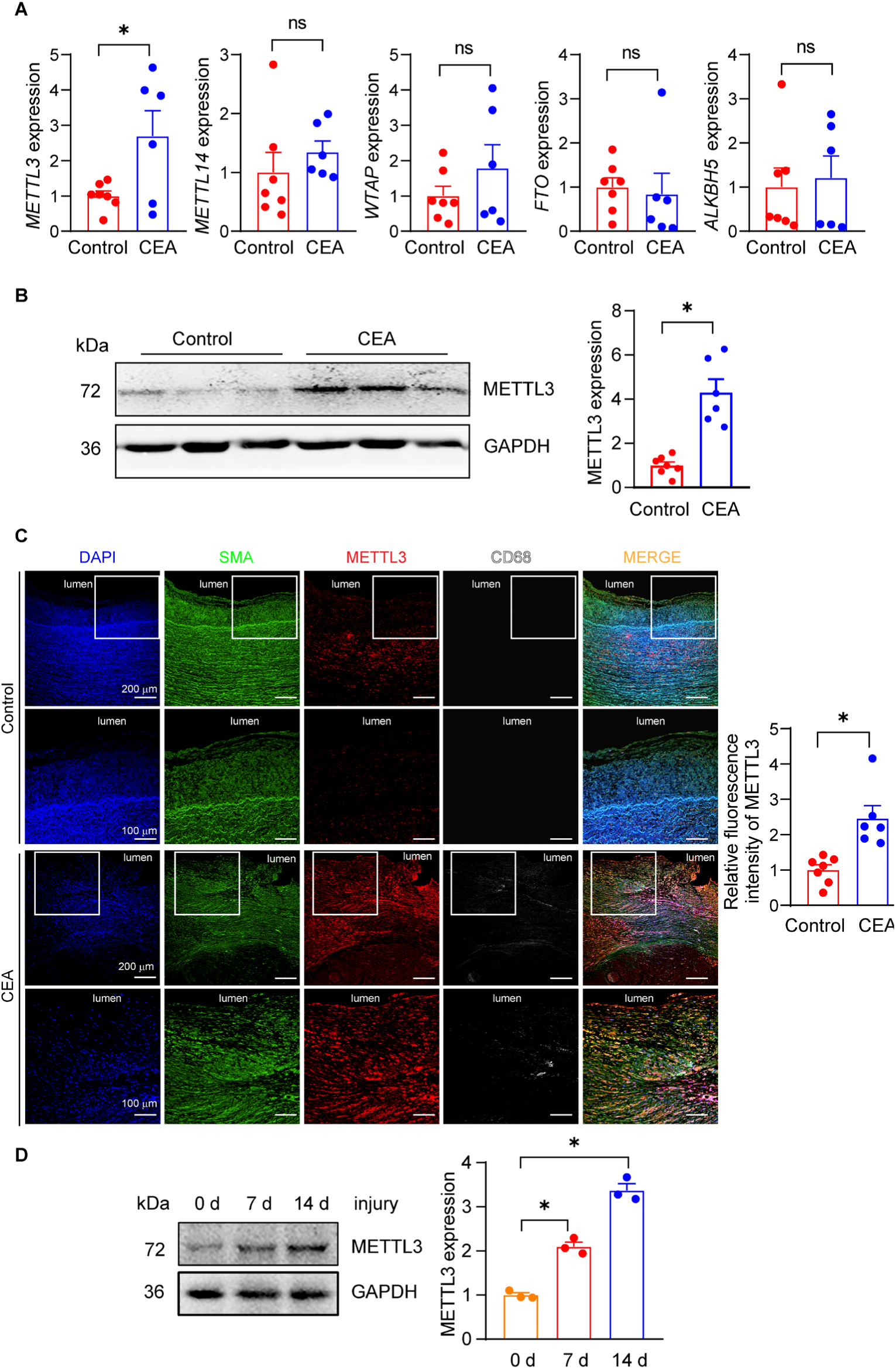
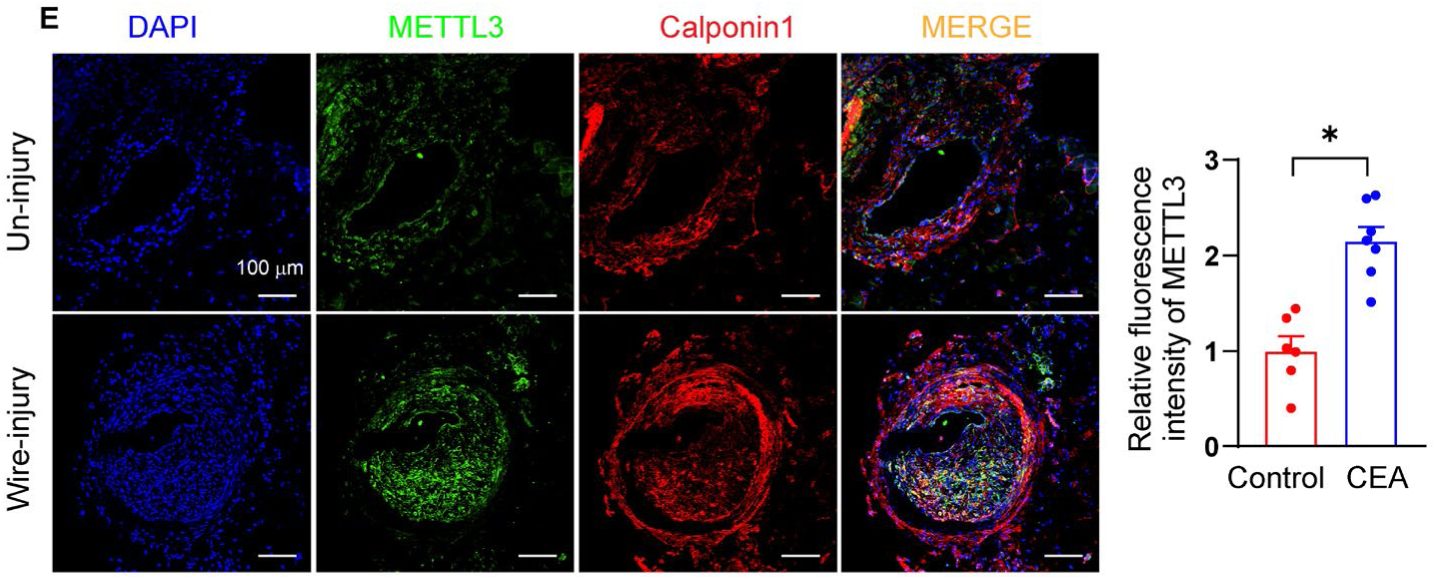
METTL3 is upregulated in mouse and human neointimal arteries. **A.** RT-qPCR analysis of the relative mRNA levels of m^6^A methyltransferases and demethyltransferases expression in human neointimal tissues collected from carotid endarterectomy (CEA) and normal carotid arteries, including methyltransferase-like 3 (METTL3), Wilms tumor 1-associated protein (WTAP), methyltransferase-like 14 (METTL14), fat mass and obesity-associated protein (FTO), and alkB homolog 5, RNA demethylase (ALKBH5). N=6-7, unpaired two-tailed Student’s *t* test. \**P*<0.05. **B.** Representative Western blotting analysis and quantification of METTL3 expression in human neointimal tissues collected from carotid endarterectomy and normal carotid arteries. N=6-7, unpaired two-tailed Student’s *t* test, \**P*<0.05. **C.** Immunofluorescence staining of METTL3 (red), Smooth Muscle Actin (SMA) (green) and CD68 (gray) in cross-sections of human neointimal tissues collected from CEA and normal carotid arteries. DAPI stained cell nucleus. N=6-7, unpaired two-tailed Student’s *t* test, \**P*<0.05. **D.** Representative Western blotting showing METTL3 expression in carotid arteries from 10-week-old male mice subjected to wire injury at 0, 7 and 14 days postinjury. GAPDH was used as an internal control. Each sample contained 4 carotid arteries, n=3 for each group. One-way ANOVA followed by Tukey’s test. \**P*<0.05. **E.** Immunofluorescence staining of METTL3 (green) and Calponin1 (red) in cross-sections of mouse carotid artery from 10-week-old male mice subjected to wire injury at 28 days postinjury. DAPI stained cell nucleus. n=6-7, unpaired two-tailed Student’s *t* test, \**P*<0.05.

### VSMC-specific knockout of METTL3 ameliorates postinjury neointima formation in mice

To confirm the role of METTL3 in mature VSMCs while excluding its potential effect on VSMC development, we generated tamoxifen-inducible METTL3 conditional knockout mice (*Mettl3*^flox/flox^; *Myh11*-CreER^T2^, named *Mettl3*^SMCKO^) (Figure 2A and Supplementary Figure 2B). Since the bacterial artificial chromosome containing *Myh11*-CreER^T2^ was inserted on the Y chromosome, only male mice were used for investigation. Tamoxifen was administered to 8-week-old *Mettl3*^SMCKO^ mice to achieve VSMC-confined depletion of METTL3 compared with age-matched 3 types of control wildtype mice (*Myh11*-CreER^T2^ with Tamoxifen injection, *Mettl3*^flox/flox^ with tamoxifen injection, and *Mettl3*^flox/flox^*; Myh11*-CreER^T2^ with corn oil injection mice) (Figure 2B and Supplementary Figure 2C). Subsequently, wire injury of the carotid arteries was induced in tamoxifen-induced *Mettl3*^SMCKO^ mice as well as 3 types of control wildtype mice, and neointimal hyperplasia was evaluated at 28 days after wire injury. VSMC-specific METTL3 deficiency did not affect mouse body weight or blood pressure (Supplementary Figure 2D) but greatly alleviated postinjury neointima formation, as evidenced by the obvious decreases in neointima area and the neointima/media ratio in *Mettl3*^SMCKO^ mice compared with 3 types of control mice (Figure 2C-D). Of interest, METTL3 expression and post-injury neointima formation were not distinguishing in these 3 types of control wildtype mice with distinct genotypes and treatments. Considering the side effects of tamoxifen and the potential Cre toxicity of *Myh11*-CreER^T2 19, 20^, *Myh11*-CreER^T2^ mice with Tamoxifen injection were applied as control wildtype mice (*Mettl3*^SMCWT^) for subsequent evaluation.

**Figure 2.**
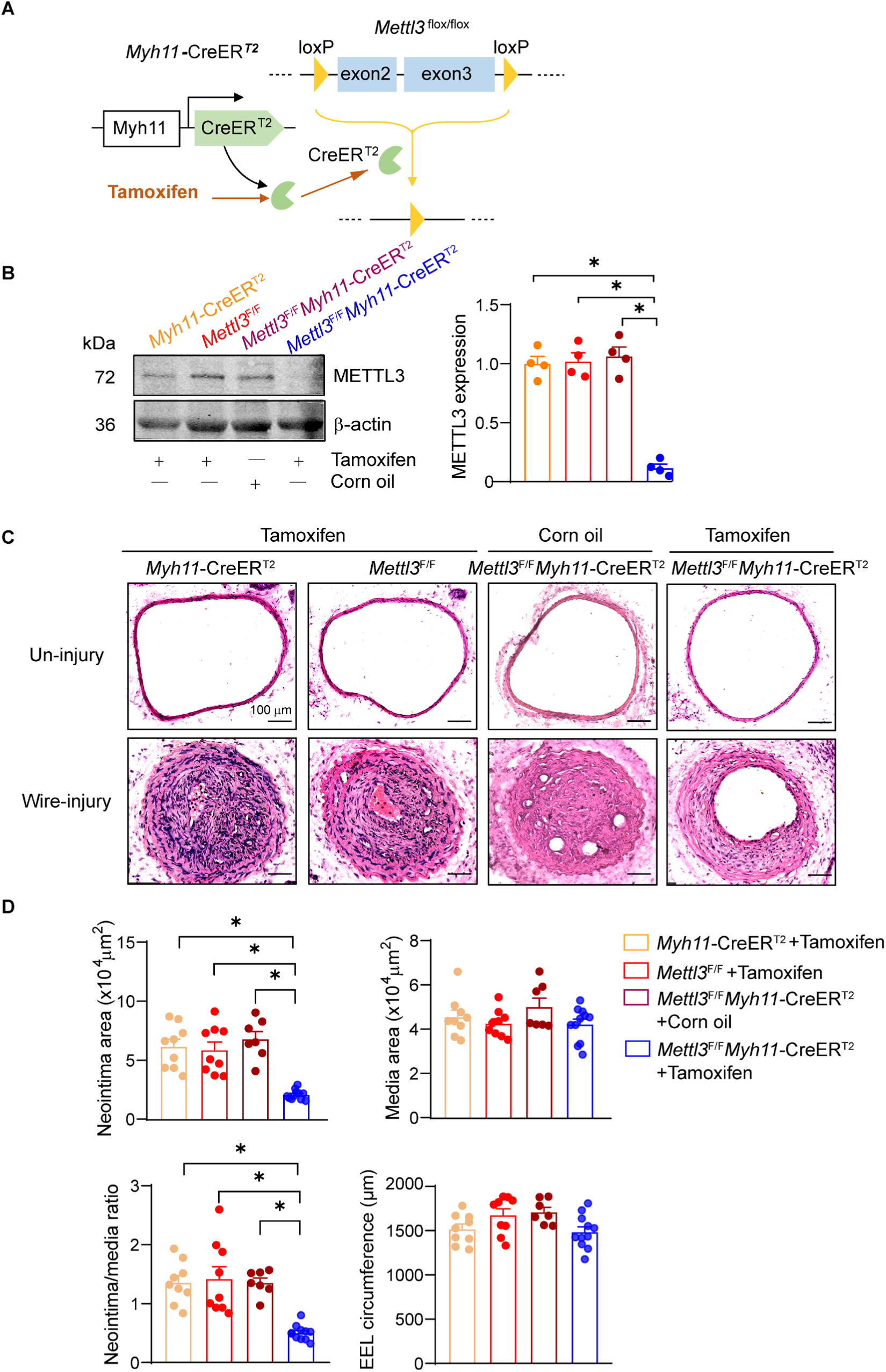
METTL3 deficiency in vascular smooth muscle cells (VSMCs) ameliorates postinjury neointima formation in mouse carotid arteries. **A.** Mouse breeding strategy for obtaining SMC-specific METTL3 knockout (*Mettl3*^flox/flox^; *Myh11*-CreER^T2^) mice. **B.** Western blotting of METTL3 expression in carotid arteries from *Myh11*CreER^T2^ with tamoxifen, *Mettl3*^flox/flox^ with tamoxifen, *Mettl3*^flox/flox^ *Myh11*CreER^T2^ with corn oil or tamoxifen treatment (75 mg/kg for 5 consecutive days) mice. GAPDH was used as an internal control. n=4, one-way ANOVA followed by Tukey’s test, *P<0.05. **C.** Hematoxylin and eosin (H&E) staining of cross-sections of sham-operated and wire-injured carotid arteries from tamoxifen-induced male *Myh11*CreER^T2^ with tamoxifen, *Mettl3*^flox/flox^ with tamoxifen, *Mettl3*^flox/flox^ *Myh11*CreER^T2^ with corn oil or tamoxifen mice 28 days after the operation. Scale bar = 100 µm. **D.** Quantitative analysis of the intima areas, media areas, neointima-to-media ratios, and external elastic lamina (EEL) circumferences in H&E-stained cross-sections. n=7-11 mice, one-way ANOVA followed by Tukey’s test, \**P*<0.05.

### METTL3 ablation represses VSMC proliferation

In view of the driving role of VSMC proliferation and migration in contributing to neointima formation, we tested whether METTL3 affects these VSMC behaviors. First, CCK-8 assays revealed that PDGF-BB or FBS induced comparable VSMC proliferation, whereas METTL3 silencing overtly suppressed the effect of PDGF-BB or FBS induction (Figure 3A). Furthermore, the EdU incorporation assays showed that there was significantly less EdU-positive staining in METTL3-silenced VSMCs than in control cells (Figure 3B), also indicating the suppression of VSMC proliferation. Since enhanced cell cycle progression is directly related to VSMC proliferation, the cell cycle analysis exhibited that the knockdown of METTL3 blocked the G1 to S phase transition in VSMCs (Supplementary Figure 3A). Meanwhile, we determined the expression of Cyclin D1 and E1, which are critical signaling molecules that promote cell cycle progression. As a result, knockdown of METTL3 significantly downregulated Cyclin D1 and E1 protein expression (Figure 3C-D), suggesting decreased cell cycle progression. These data collectively suggested that VSMC proliferation was suppressed by METTL3 knockdown. Moreover, we also measured PDGF-BB-induced VSMC migration via both trans-well migration and wound healing assays and found that METTL3 silencing did not influence VSMC migration (Supplementary Figure 3B-C), in line with no impact on migration-related gene expression in VSMCs by METTL3 knockdown (Supplementary Figure 3D). Of note, VSMC apoptosis has been reported also associated with cell quantity and neointima formation ^21^. We accordingly performed TUNEL staining and found that METTL3 knockdown did not affect VSMC apoptosis (Supplementary Figure 3E), excluding the potential impact of METTL3 on VSMC apoptosis.

**Figure 3.**
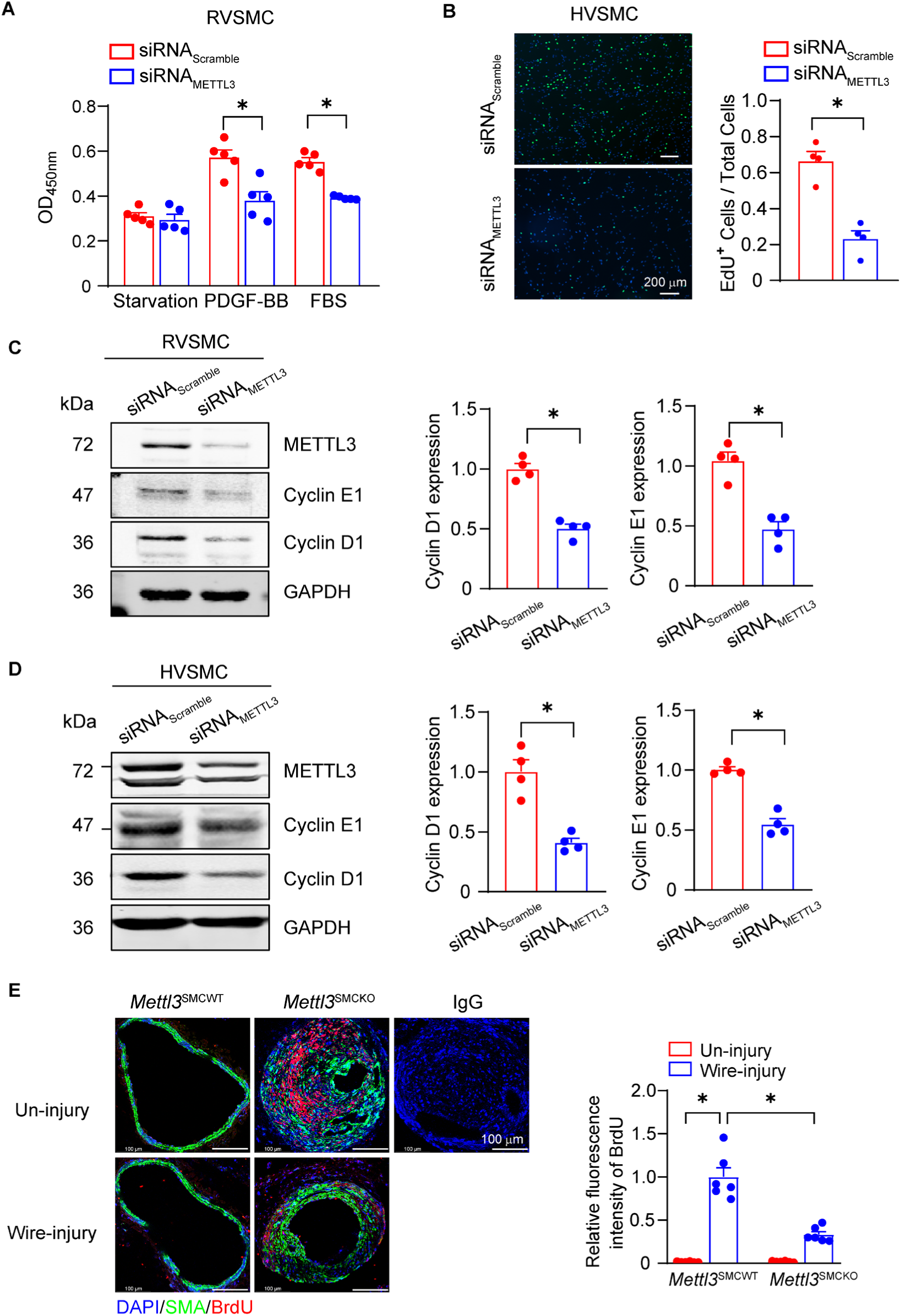
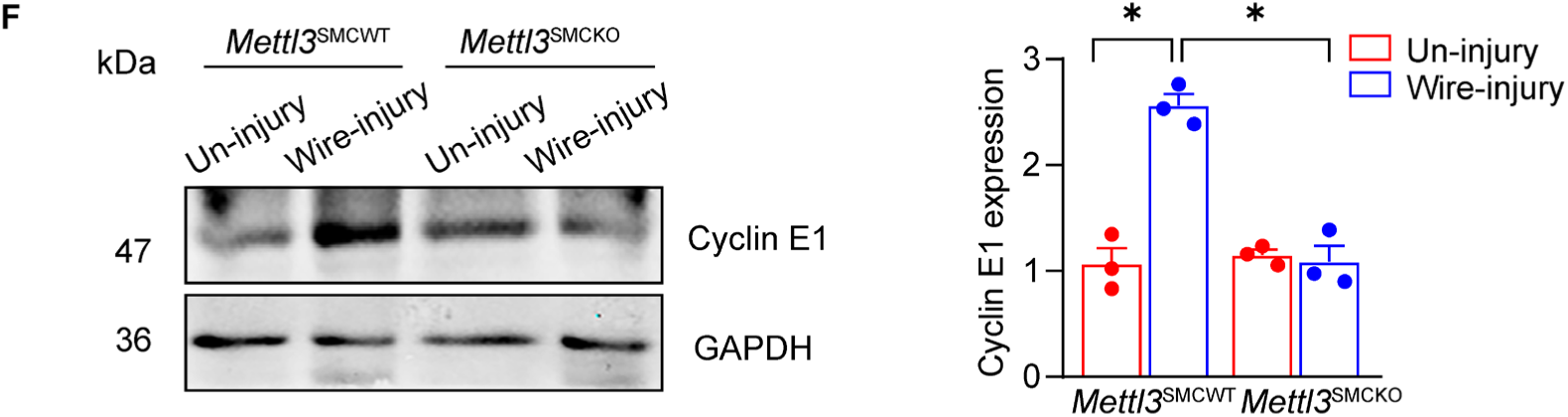
METTL3 deficiency inhibits VSMC proliferation. **A.** Cell counting kit-8 (CCK-8) proliferation assays of primary rat VSMCs (RVSMC) transfected with scramble siRNA or METTL3 siRNA (20 nM) followed by PDGF-BB (platelet derived growth factor BB) or 10% FBS (fetal bovine serum) treatment in serum-free culture media for 24 hours. n=5, two-way ANOVA followed by Tukey’s test, \**P*<0.05. **B.** EdU (5-Ethynyl-2’-deoxyuridine) incorporation assays of primary human VSMCs (HVSMC) transfected with scramble siRNA or METTL3 siRNA (20 nM) in serum-containing culture media. Scale bar = 200 µm. n=4, one-way ANOVA followed by Tukey’s test, \**P*<0.05. **C.** Representative Western blotting analysis and quantification of Cyclin D1 and Cyclin E1 expression in primary rat VSMCs transfected with scrambled siRNA or METTL3 siRNA (20 nM) in serum-containing media. n=4, unpaired two-tailed Student’s t test, \**P*<0.05. **D.** Representative Western blotting analysis and quantification of Cyclin D1 and Cyclin E1 expression in primary human VSMCs transfected with scrambled siRNA or METTL3 siRNA (20 nM) in serum-containing media. n=4, unpaired two-tailed Student’s t test, \**P*<0.05. **E.** Immunofluorescence staining of BrdU (5-bromodeoxyuridine) incorporation in the carotid arteries of 12-week-old *Mettl3* ^SMCWT^ and *Mettl3*^SMCKO^ mice at 28 days post wire injury. Scale bar = 50 μm. The data from each mouse were collected from 12 random high power fields (HPFs). n=6 mice, one-way ANOVA followed by Tukey’s test, \**P*<0.05. **F.** Representative Western blotting analysis and quantification of Cyclin E1 expression in the carotid arteries of 12-week-old *Mettl3* ^SMCWT^ and *Mettl3*^SMCKO^ mice at 14 days post wire injury. Each sample contained 4 carotid arteries. n=3 for each group, one-way ANOVA followed by Tukey’s test, \**P*<0.05.

Next, we validated the effects of METTL3 on VSMCs *in vivo*. BrdU incorporation assays in wire-injured mice displayed *Mettl3*^SMCKO^ mice exhibited fewer BrdU^+^ VSMCs in wire-injured carotid arteries compared with control *Mettl3*^SMCWT^ mice (Figure 3E). Moreover, Western blotting revealed that the upregulation of Cyclin E1 in wire-injured carotid arteries was suppressed by METTL3 deficiency in VSMCs (Figure 3F). Nevertheless, TUNEL staining did not exhibited substantial VSMC apoptosis in neointima area, suggesting that METTL3 deficiency did not enhance VSMC apoptosis (Supplementary Figure 3F). These *in vivo* data further supported that METTL3 mainly regulated VSMC proliferation rather than apoptosis related to neointima formation. Therefore, METTL3 deficiency inhibited VSMC proliferation without the impacts on migration and apoptosis.

### METTL3 methylates SGK1 mRNA and upregulates its expression in VSMCs

To elucidate the mechanism underlying the pro-proliferative effect of METTL3 on VSMCs, we simultaneously performed m^6^A methylated RNA immunoprecipitation sequencing (MeRIP-seq) and bulk RNA sequencing (RNA-seq) in VSMCs with or without METTL3 overexpression (Figure 4A, Supplementary Figure 4 and Supplementary Dataset I and II). We identified 135 genes with increased mRNA m^6^A modifications and altered mRNA expression upon METTL3 overexpression through combined MeRIP-seq and bulk RNA-seq analyses (Supplementary Dataset III). Among these 135 genes, we identified 5 candidates involved in VSMC proliferation or neointimal hyperplasia ^15, 22–25^ as potential targets of METTL3-mediated m^6^A modification that possibly mediate the effect of METTL3 on neointima formation. Next, we performed quantitative PCR (qPCR) to validate that SGK1 was most significantly upregulated upon METTL3 overexpression (Figure 4B and Supplementary Figure 5A), in line with the RNA-seq data. Conversely, METTL3 silencing downregulated SGK1 at both the mRNA and protein levels in VSMCs (Figure 4C-D). In addition, SGK1 was substantially upregulated in the neointima of wire-injured carotid arteries from mice, but downregulated by VSMC specific METTL3 deficiency (Figure 4E-F and). Additionally, SGK1 was upregulated in the neointima of patients who underwent carotid endarterectomy (Supplementary Figure 5B-C), in parallel with the upregulation of METTL3 shown in Figure 1A-C. Taken together, these data suggested that METTL3 upregulated SGK1 expression in VSMCs.

**Figure 4.**
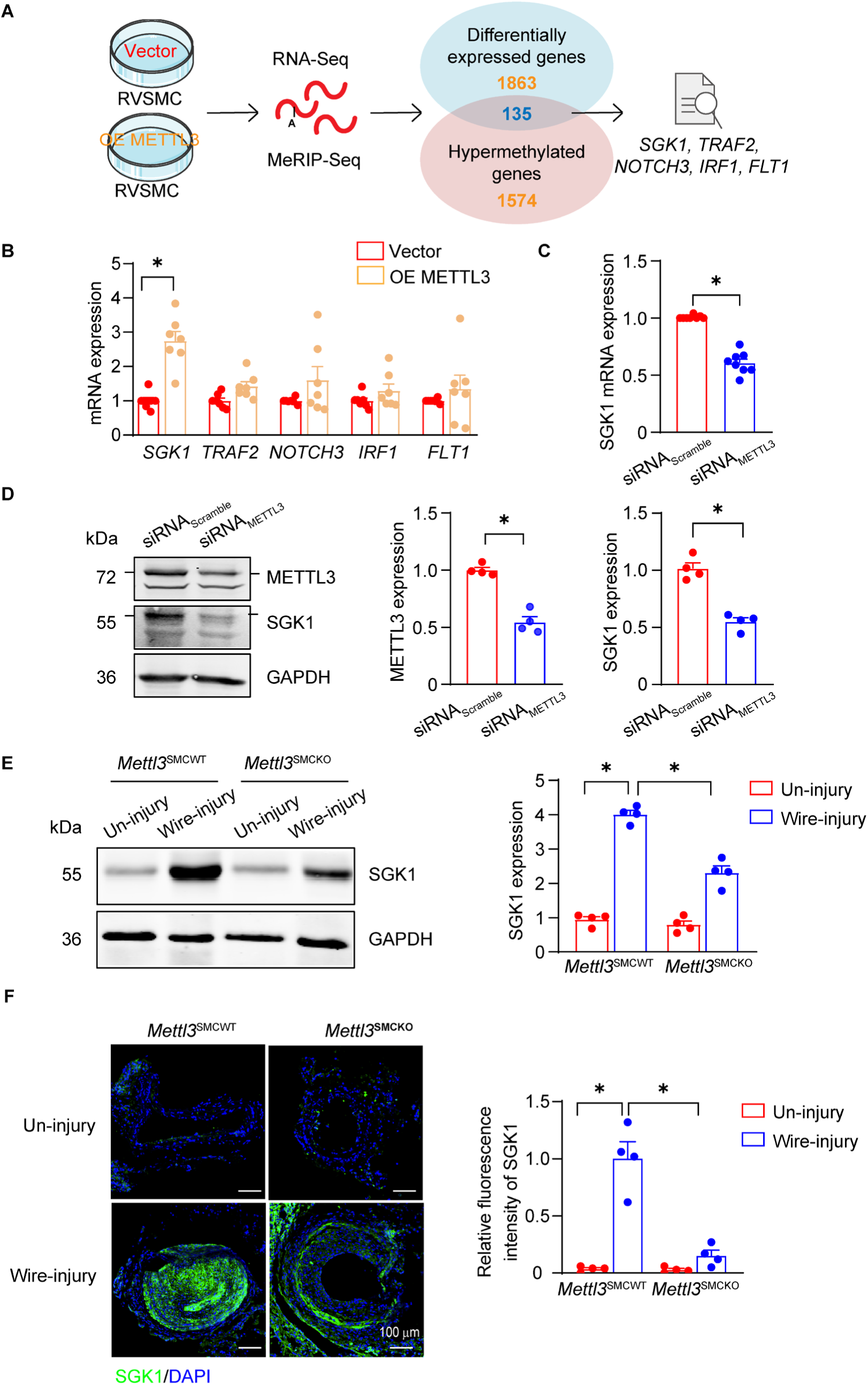
METTL3 regulates SGK1 m^6^A modification and expression in VSMCs. **A.** Schematic illustration of MeRIP (m^6^A-specific methylated RNA immunoprecipitation)-seq and RNA-seq of primary rat VSMCs transfected with the METTL3 plasmid or vector plasmid for 48 hours. The Venn diagram shows the potential targets that were hypermethylated according to MeRIP-seq and differentially expressed according to RNA-seq, including SGK1 (Serum/Glucocorticoid Regulated Kinase 1), TRAF2 (TNF Receptor Associated Factor 2), NOTCH3 (Neurogenic Locus Notch Homolog Protein 3), IRF1 (Interferon Regulatory Factor 1) and FLT1 (Fms Related Receptor Tyrosine Kinase 1). **B.** RT‒qPCR analysis of primary rat VSMCs transfected with the METTL3 plasmid or vector plasmid for 48 hours. n=7, unpaired two-tailed Student’s *t* test, \**P*<0.05. **C.** RT‒qPCR analysis of SGK1 mRNA expression in primary rat VSMCs transfected with scrambled siRNA (20 nM) or METTL3 siRNA (20 nM) in serum-containing media for 48 hours. n=8, unpaired two-tailed Student’s *t* test, \**P*<0.05. **D.** Representative Western blotting analysis and quantification of METTL3 and SGK1 expression in primary rat VSMCs transfected with scrambled siRNA (20 nM) or METTL3 siRNA (20 nM) in serum-containing media for 48 hours. n=4, unpaired two-tailed Student’s *t* test, \**P*<0.05. **E.** Representative Western blotting analysis and quantification of SGK1 expression in carotid arteries from sham-operated or wire-injured 10-week-old male mice at 7 days post-operation. GAPDH was used as an internal control. Each sample contained 4 carotid arteries. n=4 for each group, two-way ANOVA followed by Tukey’s test, \**P*<0.05. **F.** Immunofluorescence staining of SGK1 (green) in cross-sections of carotid arteries from sham-operated or wire-injured 10-week-old male mice at 28 days post-operation. Scale bar=100 μm. DAPI stained cell nucleus. The data for each mouse were collected from 12 random HPFs. n=4 mice, two-way ANOVA followed by Tukey’s test, \**P*<0.05.

To substantiate the methylating effect of METTL3 on SGK1 mRNA, we initially performed the analysis on peak calling of the MeRIP-seq data. Consequently, we found that differentially enriched m^6^A peaks in the rat SGK1 transcript resided mainly within the scope of exon 1, spanning from the 5’-untranslated region (5’UTR) to the starting region of the coding region. In light of this, qPCR primers were designed accordingly to examine m^6^A antibody-mediated immunoprecipitation of methylated SGK1 transcripts in rat VSMCs. As displayed in Figure 5A, METTL3 silencing significantly decreased the m^6^A enrichment of SGK1 transcripts, validating the methylation of SGK1 mRNA by METTL3. Considering the possible peak shift bias in MeRIP sequencing and the distribution of the m^6^A motif RRACH (R = A or G; H = A, C, or U) ^26^ in the preceding part of the rat SGK1 transcript, an extension of ∼ 230 nucleotides was included in the candidate methylation region (1-367 nucleotides in the 5’ end of the transcript) to investigate the methylation sites. Then, MeRIP-qPCR with primers allowing amplification of distinct truncations of the candidate region was performed (Figure 5B). Our data indicated that rat SGK1 mRNA transcripts ranging from 1 to 136 nucleotides in length as well as from 114 to 257 nucleotides in length were successfully enriched by m^6^A antibodies, whereas those ranging from 236 to 367 nucleotides in length were not enriched (Figure 5C). Subsequently, we inserted a transcript of 257 nucleotides, including 4 RRACH motifs, into the pGL3-promoter vector and generated 4 additional constructs bearing site mutations corresponding to distinct RRACH motifs (Figure 5D). These wild-type (SGK1-wt) or mutant (SGK1-mut) constructs were then individually transfected into A7r5 smooth muscle cell line. To accurately assess the methylation levels of above four chimeric RNA exogenously introduced from these constructs, we performed MeRIP-qPCR analysis and the results suggested that mutations in the third (A147U) or fourth (A210U) motif, rather than the first (A39U) or second (A72U) motif, significantly impaired m^6^A antibody-mediated enrichment of the chimeric transcript (Figure 5E). Thus, both A147 and A210 in rat SGK1 mRNA are thought to be methylated by METTL3. To confirm whether these two m^6^A motifs in rat SGK1 transcripts are conserved between mice and humans, we performed bioinformatics analysis of cross-species SGK1 mRNA sequence alignment. Consequently, the homology of the A147 m^6^A motif is limited in mouse and rat SGK1 transcripts, whereas the A210 site is highly conserved across the three species (Supplementary Figure 6), suggesting that similar site-specific m^6^A modifications probably also exist in mouse and human SGK1 transcripts. Taken together, these findings demonstrated that METTL3 directly methylated SGK1 mRNA and promoted its expression in VSMCs.

**Figure 5.**
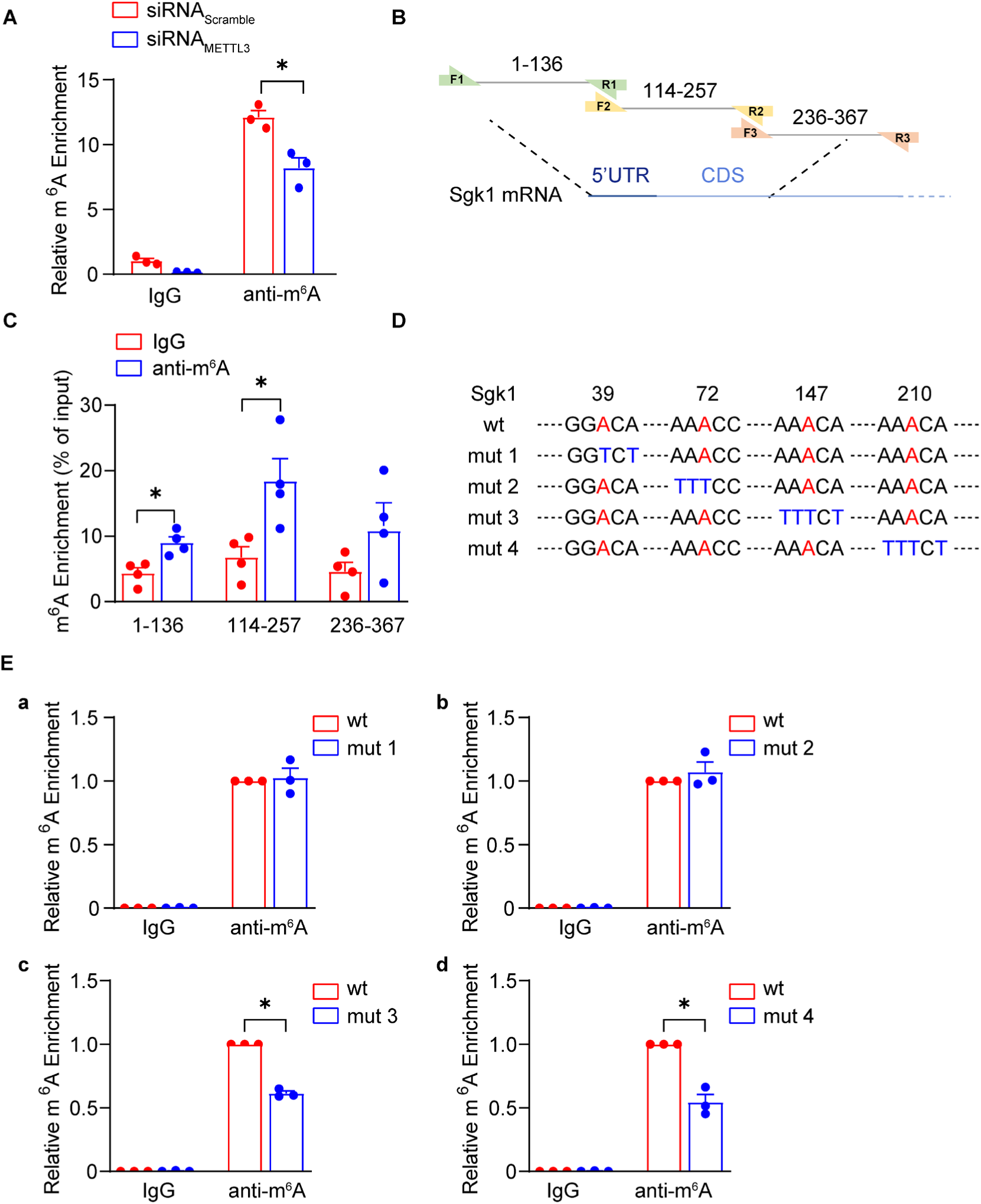
METTL3 mediates m^6^A modification downstream of the transcription start site in the SGK1 mRNA transcript. **A.** MeRIP-qPCR analysis of SGK1 mRNA m^6^A modification in primary rat VSMCs transfected with scrambled siRNA (20 nM) or METTL3 siRNA (20 nM) for 48 hours. Rabbit IgG was used as a negative control. n=3, two-way ANOVA followed by Tukey’s test, \**P*<0.05. **B.** Schematic of the potential m^6^A modification regions in the rat SGK1 mRNA transcript. **C.** MeRIP-qPCR analysis of primary rat VSMCs using primers targeting the distinct potential m^6^A methylation regions of the SGK1 mRNA transcript. n=3, unpaired two-tailed Student’s *t* test, \**P*<0.05. **D.** Schematics of the predicted position of m^6^A motifs (wt) and the individual mutations in different m^6^A motifs (mut 1-4) within the m^6^A modification region of the rat SGK1 mRNA transcript. **E.** MeRIP-qPCR analysis of primary rat VSMCs transfected with pGL3-promoter vectors encoding SGK1 wt or SGK1 mut 1-4 transcripts. Rabbit IgG was used as a negative control. n=3, two-way ANOVA followed by Tukey’s test, \**P*<0.05.

### m^6^A modification of SGK1 mRNA facilitates its transcription in a YTHDC1–dependent manner in VSMCs

Next, we explored how METTL3-mediated m^6^A modification upregulated SGK1 mRNA expression. Accordingly, SGK1 mRNA stability and precursor messenger RNA (pre-mRNA) levels were evaluated. The results showed that METTL3 knockdown did not affect the stability of SGK1 mRNA (Figure 6A) but markedly decreased the abundance of SGK1 pre-mRNA in VSMCs (Figure 6B). Since pre-mRNA is directly transcribed from the genome DNA inside the nucleus, these results suggested that METTL3-mediated m^6^A modification likely increases SGK1 gene transcription. RNA m^6^A modification dictates mRNA metabolism mainly via m^6^A readers, we next explored if certain critical reader proteins contribute to m^6^A modification-regulated SGK1 gene transcription. YTHDC1, as a well-recognized m^6^A reader, has recently been reported to enhance the transcription of m^6^A-modified mRNAs ^5^. Therefore, we tested if YTHDC1 affects SGK1 gene transcription and expression. Similar to METTL3, YTHDC1 knockdown downregulated SGK1 expression at both the mRNA and protein levels in VSMCs (Figure 6C-D and Supplementary Figure 7A). Moreover, YTHDC1 silencing inhibited PDGF-BB-induced VSMC proliferation and accordingly downregulated Cyclin D1 and E1 expression in VSMCs (Figure 6E-F), whereas overexpression of SGK1 by adenovirus infection efficiently reversed YTHDC1 silencing-suppressed VSMC proliferation (Figure 6G and Supplementary Figure 7B), indicating that YTHDC1 is also functionally involved in VSMC proliferation through modulating SGK1 expression.

**Figure 6.**
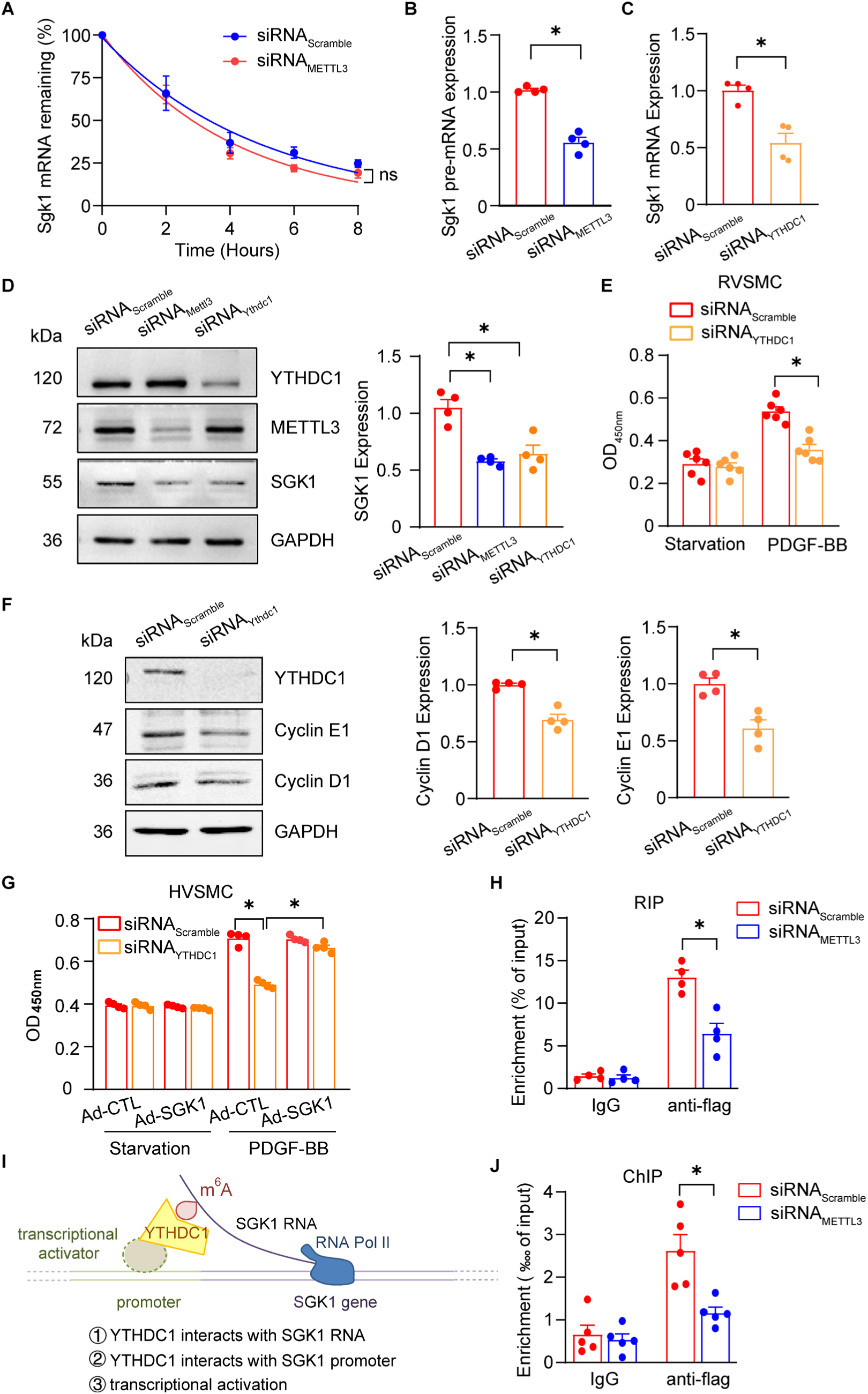
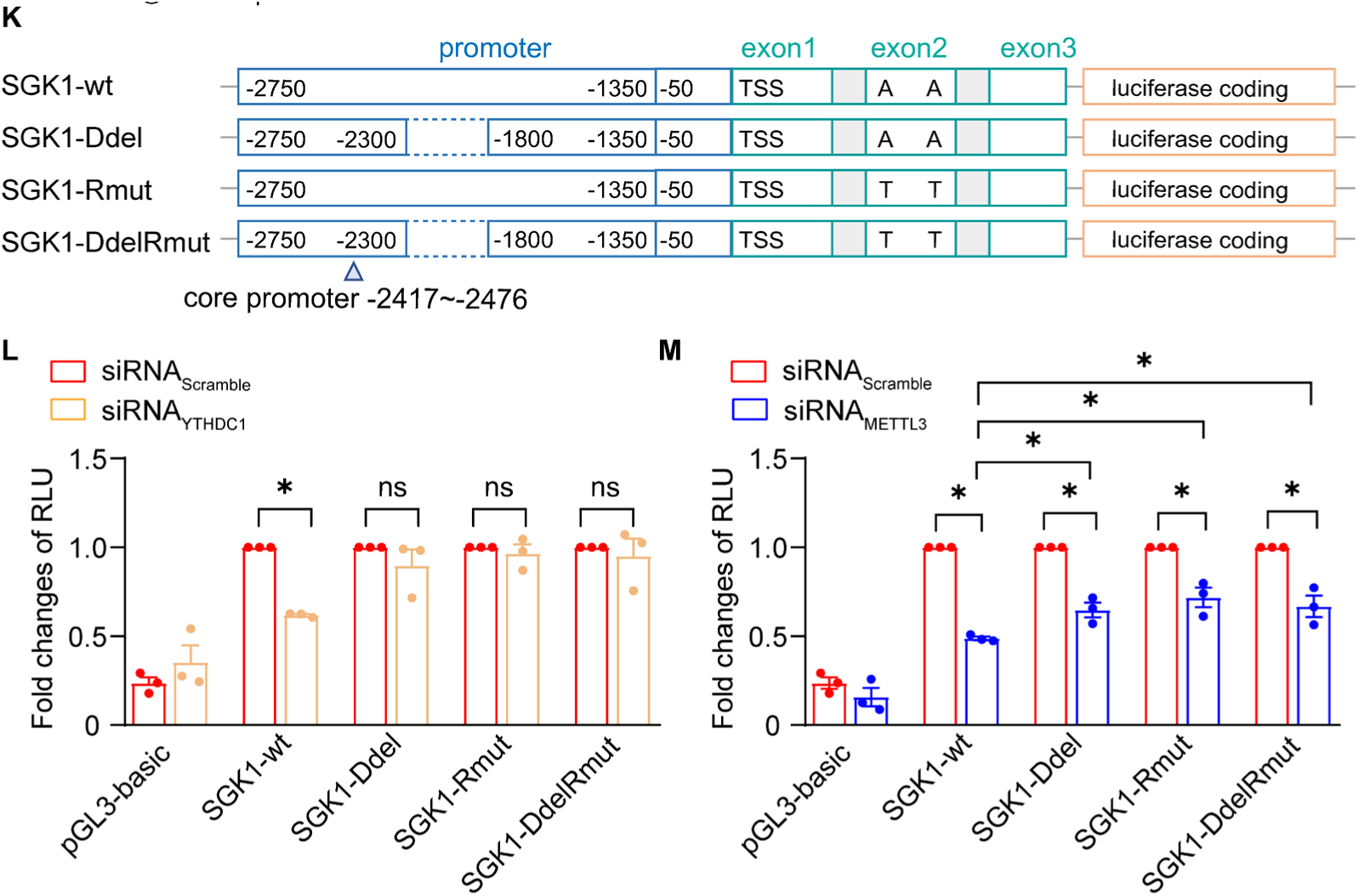
YTHDC1 promotes SGK1 gene transcription through m^6^A modification of the SGK1 transcript. **A.** RT‒qPCR analysis of the remaining SGK1 mRNA in scramble siRNA or METTL3 siRNA (20 nM)-transfected rat primary VSMCs treated with actinomycin D (2 μg/mL) and harvested at 0-, 2-, 4-, 6-, and 8-hour time points following actinomycin D addition. The half-life (t_1/2_) of SGK1 mRNA was fitted and calculated independently for each experiment. n=4, unpaired two-tailed Student’s *t* test, ns, no significance. **B.** RT‒qPCR analysis of SGK1 pre-mRNA expression in primary rat VSMCs transfected with scrambled siRNA or METTL3 siRNA for 48 hours in serum-containing media. n=4, unpaired two-tailed Student’s *t* test. \**P*<0.05. **C.** RT‒ qPCR analysis of SGK1 expression in primary rat VSMCs transfected with scrambled siRNA or YTHDC1 siRNA for 48 hours in serum-containing media. n=4, unpaired two-tailed Student’s *t* test. \**P*<0.05. **D.** Representative Western blotting analysis and quantification of METTL3, YTHDC1 (YTH domain-containing protein 1) and SGK1 protein expression in A7r5 cells transfected with scrambled siRNA, METTL3 siRNA or YTHDC1 siRNA for 48 hours in serum-containing media. n=4, one-way ANOVA followed by Tukey’s test, \**P*<0.05. **E.** CCK-8 proliferation assays of primary human VSMCs transfected with scramble siRNA or YTHDC1 siRNA (20 nM) for 48 hours followed by treatment with PDGF-BB (20 ng/mL) in serum-free media. n=6, two-way ANOVA followed by Tukey’s test, \**P*<0.05. **F.** Representative Western blotting analysis and quantification of Cyclin E1 expression in primary rat VSMCs transfected with scrambled siRNA or YTHDC1 siRNA (20 nM) for 48 hours in serum-containing media. n=4, unpaired two-tailed Student’s *t* test, \**P*<0.05. **G.** CCK-8 proliferation assays of primary human VSMCs transfected with scramble siRNA (20 nM) or YTHDC1 siRNA (20 nM) for 48 hours followed by infection with control adenovirus (Ad-CTL) or SGK1 adenovirus (Ad-SGK1) for 24 hours. PDGF-BB (20 ng/mL) was applied to induce proliferation in serum-free media for 24 hours. Two-way ANOVA followed by Tukey’s test, \**P*<0.05. **H.** RIP-qPCR analysis of A7r5 cells stably transfected with pcDNA3.0-YTHDC1-Flag and scrambled siRNA with or without siRNA-mediated METTL3 silencing. Mouse IgG was used as a negative control. n=4, two-way ANOVA followed by Tukey’s test, \**P*<0.05. **I.** Schematic of the hypothesis that YTHDC1 promotes the transcription of SGK1. **J.** ChIP (chromatin immunoprecipitation) ‒qPCR analysis of A7r5 cells stably transfected with pcDNA3.0-YTHDC1-Flag and scrambled siRNA with or without siRNA-mediated METTL3 silencing. Mouse IgG was used as a negative control. n=5, two-way ANOVA followed by Tukey’s test, \**P*<0.05. **K.** Schematic of the construction of pGL3 basic reporter plasmids encoding the SGK1-wt, SGK1-Ddel, SGK1-Rmut or SGK1-DdelRmut sequences for dual-luciferase reporter assays. **L.** Dual-luciferase assays in A7r5 cells transfected with scramble siRNA or YTHDC1 siRNA as well as pGL3 reporter plasmids and Renilla reporter plasmid. n=3, two-way ANOVA followed by Tukey’s test, \**P*<0.05. **M.** Dual-luciferase assays in A7r5 cells transfected with scramble siRNA or METTL3 siRNA as well as pGL3 reporter plasmids and Renilla reporter plasmid. n=3, two-way ANOVA followed by Tukey’s test, \**P*<0.05.

Furthermore, RNA immunoprecipitation assays were performed in the A7r5 smooth muscle cell line stably expressing YTHDC1-Flag to examine the association of YTHDC1 with m^6^A-modified SGK1 mRNA. As expected, the SGK1 transcript bearing 1-257 nucleotides was obviously enriched in YTHDC1-mediated immunoprecipitates, but METTL3 knockdown or SGK1 mRNA m^6^A motif mutation sharply decreased this enrichment (Figure 6H and Supplementary Figure 8A-D), demonstrating that YTHDC1 recognized METTL3-mediated m^6^A modifications on SGK1 mRNA. Recent studies have shown that the mechanism of YTHDC1-regulated gene transcription involves the use of a scaffold that links the nascent m^6^A-decorated transcript and its promoter DNA via the recognition of m^6^A modifications and its association with transcriptional activators, respectively ^6, 27^. Notably, the m^6^A sites of SGK1 mRNA span the proximity of the gene transcription start site (TSS) as well as the promoter region. Hence, we proposed a similar working model of YTHDC1 involved in SGK1 gene transcription (Figure 6I). Chromatin immunoprecipitation combined with qPCR (ChIP‒qPCR) assays were performed to explore the interaction of YTHDC1 with SGK1 promoter DNA. According to a previous study ^28^, serial pairs of primers were designed to amplify distinct SGK1 promoter segments and confirm the binding sites of YTHDC1. The mapping results suggested that YTHDC1 was predominantly associated with the promoter region ranging from 1800 bp to 2300 bp upstream of the SGK1 TSS (Supplementary Figure 8E-F), whereas knockdown of METTL3 significantly decreased the binding of YTHDC1 to the SGK1 promoter (Figure 6J). These data confirmed that YTHDC1 functions as a scaffold dependent on METTL3-mediated m^6^A modification to link the SGK1 transcript to its promoter DNA.

To verify whether the scaffold role of YTHDC1 regulates SGK1 transcription, a set of chimeric reporter constructs (pGL3) simultaneously harboring the SGK1 promoter with or without the deletion of the YTHDC1 binding sequence (Ddel) and a gene sequence involving mRNA methylation regions (1-257) with or without m^6^A site (A147 and A210) mutations (Rmut) were generated, as shown in Figure 6K. Dual luciferase activity was determined after transfecting these constructs into A7r5 smooth muscle cell line with or without YTHDC1 siRNA. Our results showed that the relative luciferase activity of cells transfected with the SGK1 native construct (pGL3-SGK1-wt) was markedly repressed by silencing YTHDC1. However, when SGK1 constructs with Ddel (pGL3-SGK1-Ddel), Rmut (pGL3-SGK1-Rmut) or both (pGL3-SGK1-DdelRmut) were used to disrupt the association between YTHDC1 and its bound promoter region or recognized methylation sites or both, the relative luciferase activities in these transfected cells were not altered by YTHDC1 knockdown (Figure 6L), indicating that YTHDC1-facilitated SGK1 expression was dependent on its association with the SGK1 m^6^A methylated transcript and promoter. Next, to confirm whether METTL3-regulated SGK1 expression was dependent on the scaffold function of YTHDC1, METTL3 siRNA was transfected into A7r5 smooth muscle cell line expressing various pGL3-SGK1 constructs. Consequently, METTL3 ablation drastically decreased the relative luciferase activity derived from cells transfected with pGL3-SGK1-wt, whereas disruption of the association between YTHDC1 and the SGK1 transcript or promoter or both significantly reversed METTL3 silencing-reduced luciferase activity (Figure 6M), suggesting that the association of YTHDC1 with m^6^A sites in the SGK1 transcript as well as the SGK1 gene promoter was involved in METTL3-facilitated SGK1 transcription. Notably, the relative luciferase activities were not reversed to the levels in the scramble siRNA-transfected cells, suggesting that other unidentified regulatory pathways of METTL3 affect SGK1 expression but are independent of the m^6^A-YTHDC1 axis.

### SGK1 overexpression reverses METTL3 deficiency-mediated suppression of VSMC proliferation and postinjury neointimal hyperplasia

To confirm whether SGK1 mediated the effect of METTL3 on VSMC proliferation, we first transduced VSMCs with adenovirus to efficiently overexpress SGK1. Consequently, SGK1 overexpression reversed the effect of METTL3 silencing on the inhibition of Cyclin E1 expression as well as proliferation in PDGF-BB-induced VSMCs (Figure 7A-C). Furthermore, an AAV9 virus with an expression cassette under the control of the SM22 promoter was utilized to induce VSMC-specific overexpression of SGK1 in control and *Mettl3*^SMCKO^ mice (Figure 7D). SGK1 overexpression in VSMCs abolished METTL3 deficiency-mediated reductions in neointima formation in mice (Figure 7E-F). These observations demonstrated that METTL3 aggravated VSMC proliferation and neointima formation by upregulating SGK1 expression.

**Figure 7.**
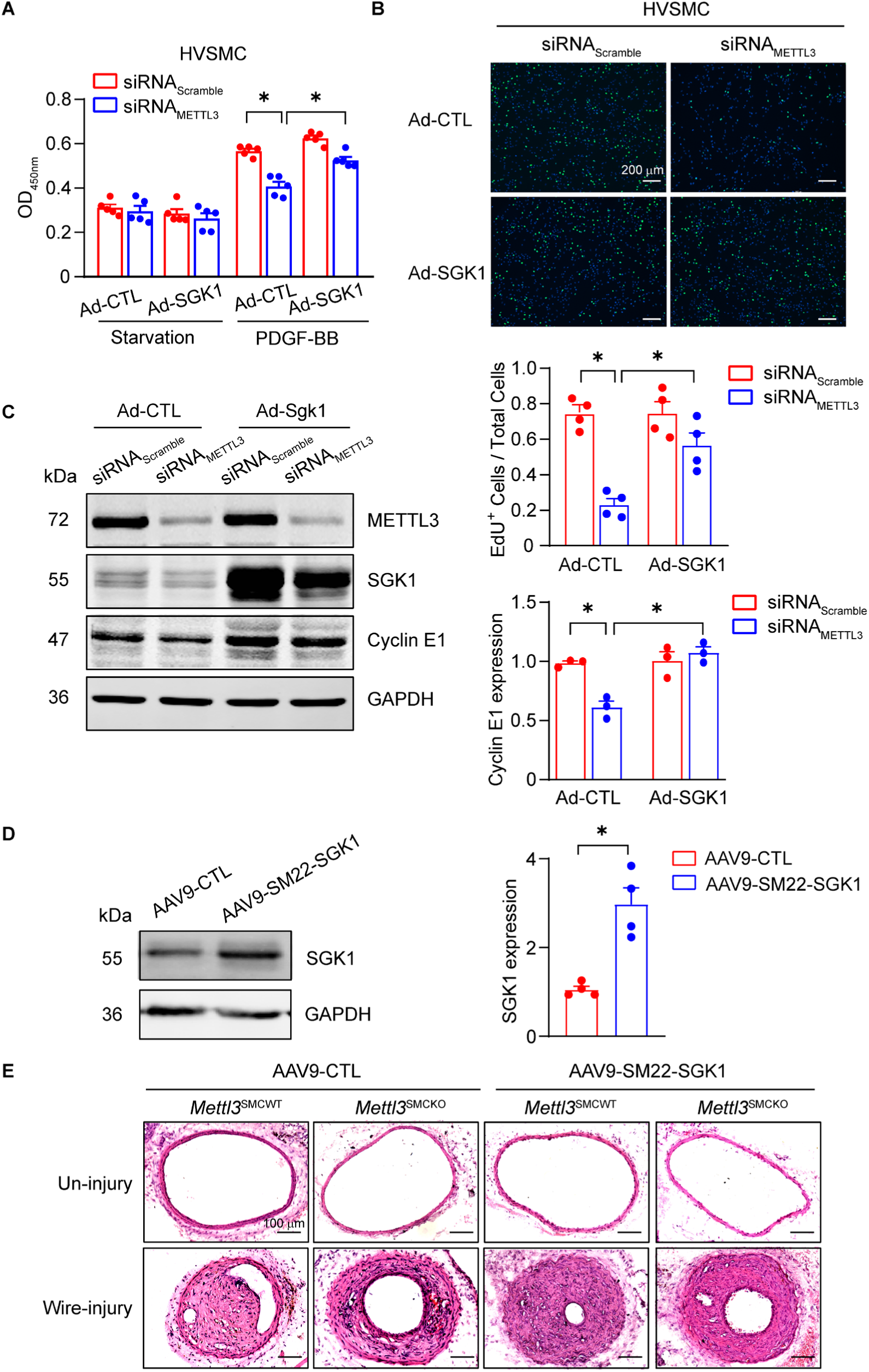
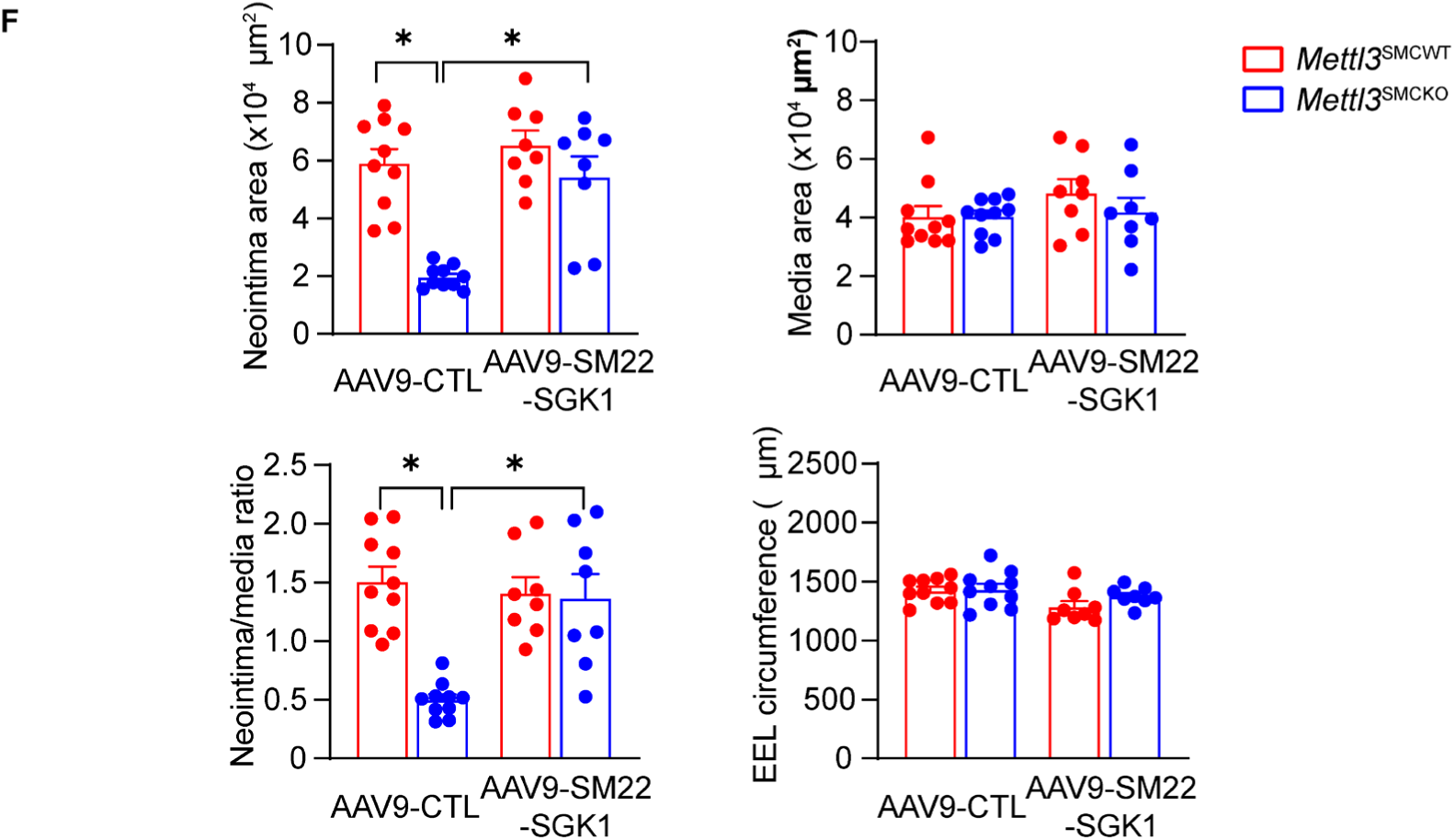
SGK1 overexpression reverses METTL3 deficiency-induced decreases in VSMC proliferation and postinjury neointima formation. **A.** CCK-8 proliferation assays of primary human VSMCs transfected with scramble siRNA (20 nM) or METTL3 siRNA (20 nM) for 48 hours followed by infection with control adenovirus (Ad-CTL) or SGK1 adenovirus (Ad-SGK1) for 24 hours. PDGF-BB (20 ng/mL) was applied to induce proliferation in serum-free media for 24 hours. Two-way ANOVA followed by Tukey’s test, \**P*<0.05. **B.** EdU incorporation assays of primary human VSMCs transfected with scramble siRNA (20 nM) or METTL3 siRNA (20 nM) for 48 hours followed by infection with control adenovirus (Ad-CTL) or SGK1 adenovirus (Ad-SGK1) for 24 hours in serum-containing media. Scale bar = 200 µm. n=4, two-way ANOVA followed by Tukey’s test, \**P*<0.05. **C.** Representative Western blotting analysis and quantification of Cyclin E1 expression in primary rat VSMCs transfected with scrambled siRNA or METTL3 siRNA for 48 hours followed by infection with control adenovirus (Ad-CTL) or SGK1 adenovirus (Ad-SGK1) for 24 hours in serum-containing media. n=3, two-way ANOVA followed by Tukey’s test, \**P*<0.05. **D.** Representative Western blotting analysis and quantification of SGK1 expression in carotid arteries from 8-week-old mice infected with AAV9 control (AAV9-CTL) or AAV9 SM22 promoter-driven SGK1 overexpression (AAV9-SM22-SGK1) virus for 14 days. GAPDH was used as an internal control. Each sample contained 4 carotid arteries. n=4, unpaired two-tailed Student’s *t* test, \**P*<0.05. **E.** Eight-week-old male *Mettl3* ^SMCWT^ and *Mettl3*^SMCKO^ mice were treated with tamoxifen (75 mg/kg) for 5 consecutive days and infected with AAV9 control (AAV9-CTL) or AAV9 SM22 promoter-driven SGK1 overexpression (AAV9-SM22-SGK1) virus for 14 days. Then, the carotid arteries of the treated and infected mice were subjected to wire injury for 28 days. H&E staining of cross-sections of sham-operated and wire-injured carotid arteries. Scale bar = 100 µm. **F.** Quantitative analysis of the intima areas, media areas, neointima-to-media ratios, and external elastic lamina (EEL) circumferences in H&E-stained cross-sections. n=8-10 mice, two-way ANOVA followed by Tukey’s test, \**P*<0.05.

## Discussion

Excessive growth of VSMCs directly contributes to the development of neointimal hyperplasia, which is related to several vascular diseases, such as the initiation of atherosclerotic plaque formation and in-stent restenosis ^29^. Investigating the regulatory mechanism of VSMC proliferation substantially deepens the understanding of the pathogenesis of neointima formation as well as provides insights for the development of novel therapeutic strategies for the treatment of related vascular diseases. In the present study, we revealed that METTL3-catalyzed m^6^A methylation of SGK1 mRNA facilitated YTHDC1-dependent SGK1 gene transcription and expression in VSMCs, subsequently promoting VSMC proliferation and aggravating neointima formation in carotid arteries after wire injury in mice (Figure 8). Thus, m^6^A RNA methylation could be a potential target for inhibiting VSMC proliferation and preventing restenosis after angioplasty and stenting.

**Figure 8.**
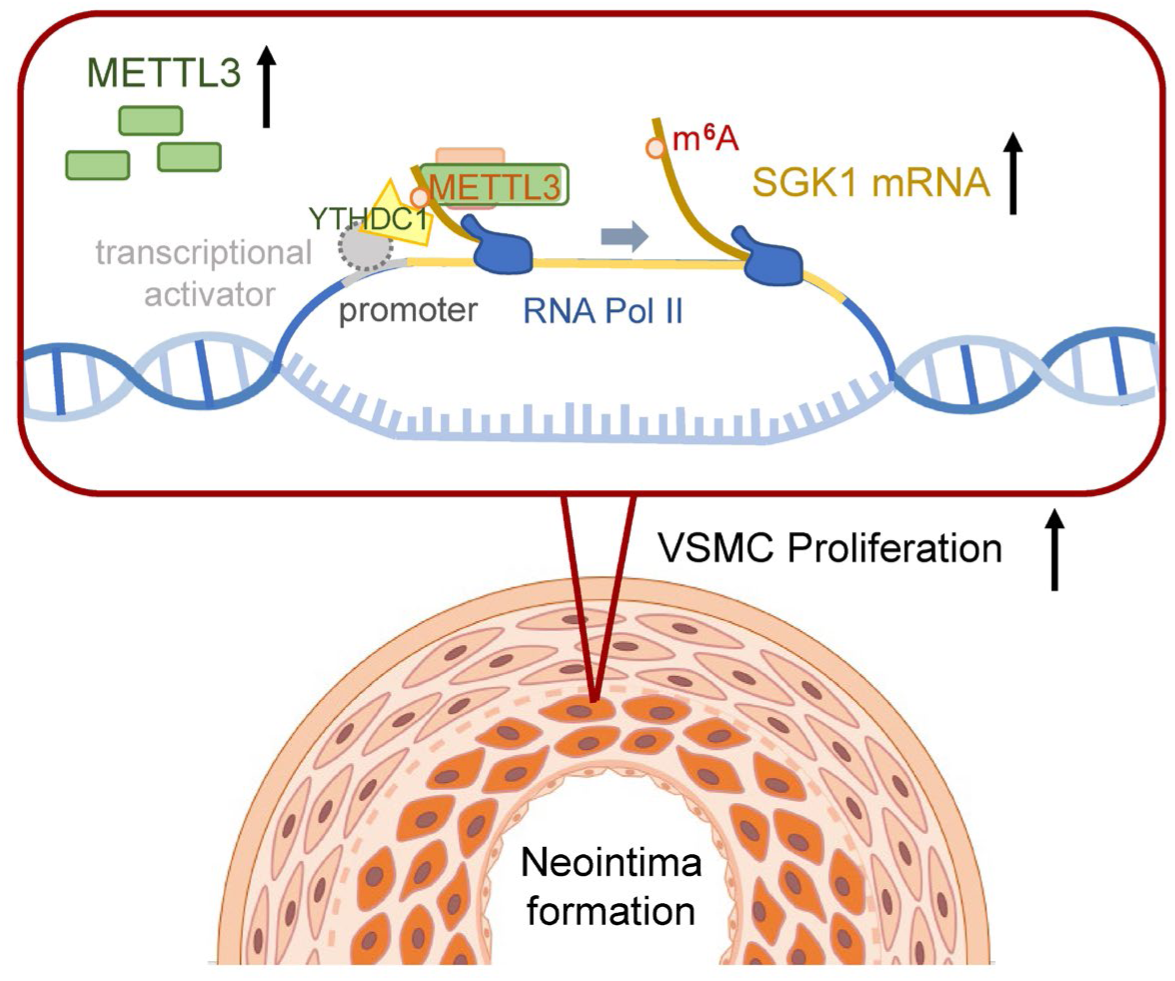
Illustration of the mechanism underlying the transcriptional regulation of SGK1 by METTL3 and YTHDC1 in neointima formation. METTL3 expression is upregulated after vascular injury, and the motifs downstream of the transcriptional start site (TSS) in nascent SGK1 transcripts are m^6^A-modified. The reader YTHDC1 can recognize and bind to nascent SGK1 transcripts containing m^6^A modifications, recruiting transcriptional activators targeting the promoter of the SGK1 gene. The synergistic effect of these molecules results in the formation of a positive feedback loop facilitating SGK1 transcription, which in turn leads to an increase in SGK1 protein expression. SGK1 promotes the proliferation and migration of VSMCs, thus aggravating neointima formation.

Epigenetic regulation plays an important role in aberrant VSMC proliferation and related vascular pathologies. The involvement of epigenetic regulation, such as microRNA and histone deacetylation, in VSMC proliferation has become evident, as preclinical studies targeting miRNAs as well as HDAC inhibitors are ongoing ^7^. The role of m^6^A RNA methylation, the most abundant RNA epigenetic modification, in regulating VSMC proliferation and neointimal hyperplasia is poorly understood. A recent *in vitro* study revealed that METTL3 overexpression inhibited serum-induced VSMC proliferation and migration, but further exploration of the transcripts targeted by m^6^A modification and validation in animal models is needed ^11^. Another study utilized direct perivascular infection with an AAV encoding shRNA to silence METTL3 specifically in the VSMCs of rat carotid arteries and found that METTL3 silencing inhibited neointimal hyperplasia in balloon-injured arteries, but the authors confusingly concluded that METTL3-mediated m^6^A methylation inhibited VSMC phenotypic switching ^10^. Overall, these limited studies remain highly controversial and lack direct evidence from VSMC-specific knockout mice. Here, we provided *in vivo* evidence using VSMC-specific METTL3 knockout mice to reveal that METTL3-catalyzed m^6^A RNA methylation promoted VSMC proliferation and aggravated neointimal hyperplasia in the carotid arteries from wire-injured mice. Interestingly, no discernible impact of METTL3-mediated m^6^A modification on VSMC migration was observed in our study, which diverges from previous *in vitro* findings. Our study broadens the understanding of the epigenetic regulation of VSMC proliferation. However, it should be notable that *Myh11*-CreER^T2^ mice we utilized here were generated by the insertion of bacterial artificial chromosome containing *Myh11*-CreER^T2^ in the Y chromosome, which exclude the application in female mice. Thus, other tool mice with VSMC-specific Cre or CreER alleles inserted in the non-Y chromosomes would be required to additional clarify the effects of METTL3 on VSMCs in female mice.

Moreover, the SGK1 transcript was identified as a specific target regulated by METTL3-catalyzed m^6^A modification in VSMCs. Previous investigations have extensively elucidated the pivotal role of SGK1 in VSMC proliferation ^22, 30^. SGK1 promoted VSMC proliferation by phosphorylating p27kip1 and modulating GSK3β-β-catenin signaling. Indeed, in a mouse vein graft model, SGK1 deficiency significantly attenuated VSMC proliferation and neointima formation. Based on these published studies, we concluded that METTL3-mediated VSMC proliferation is dependent on SGK1, which was further supported by our finding that SGK1 overexpression abolished METTL3 deficiency-inhibited VSMC proliferation and neointima formation. Although the facilitation of VSMC proliferation by SGK1 has been widely acknowledged, there has been limited investigation into the regulation of SGK1 expression in VSMCs. Growth factors (e.g., PDGF-BB, IGF-1, and BMP), mechanical stress and wire injury stimuli significantly were found to upregulate SGK1 in VSMCs, and this upregulation was dependent on MEK1, mTORC and/or SMAD signaling ^31–33^. Here, we discovered that METTL3-mediated RNA methylation plays a crucial role in the epigenetic upregulation of SGK1 expression upon exposure to pro-proliferative stimuli. Notably, our MeRIP-seq analysis identified 135 target transcripts modified by m^6^A modification with subsequent and significant changes at mRNA levels. Among them, SGK1 has been reported involved in VSMC proliferation and exhibited the most substantial upregulation of mRNA and protein expression by METTL3 overexpression. It would be rational to preferentially validate whether SGK1 mediated METTL3-modulated VSMC proliferation. Of interest, SGK1 overexpression efficiently but not completely reversed METTL3 silencing/deficiency-mediated suppression of VSMC proliferation and postinjury neointimal hyperplasia both in vitro and in vivo. This incomplete reversal of the suppressive effects mediated by METTL3 silencing/deficiency suggested additional methylated targets might be also involved in the regulation of METTL3 on VSMC proliferation, which requires further investigation. In addition to proliferation, whether METTL3 affected other behaviors or functions of VSMCs under distinct pathological conditions still need additional exploration as well, possibly involving other potential targets.

The involvement of RNA m^6^A modification in transcription has recently been revealed, but this modification has not been extensively investigated compared to its other roles in posttranscriptional regulation, such as mRNA stability, splicing, and translation ^3^. In the present study, METTL3-mediated m^6^A modification facilitated SGK1 gene transcription by recruiting the m^6^A “reader” YTHDC1. Notably, YTHDC1, among the members of the YTH family, is unique because it is localized within the nucleus ^34^. Thus, YTHDC1 can directly interact with various nuclear components, including transcriptional activators, to regulate transcription ^35^. In this context, a growing number of transcription cofactors (e.g., KDM3B, RNA polymerase II and BRD4) have been found to associate with YTHDC1 and thereby link nascent m^6^A-modified transcripts with promoter DNA ^5^. Herein, we demonstrated the dependence of YTHDC1 on METTL3-mediated m^6^A modification as a scaffold for connecting the SGK1 transcript to its promoter DNA and thus facilitating SGK1 transcription. However, the exact transcriptional activators recruited by YTHDC1 targeting SGK1 promoter DNA remain to be elucidated. In addition, YTHDC1 can promote gene transcription by engaging other mechanisms, such as affecting heterochromatin, caRNA decay and nuclear condensation ^5^. Whether these mechanisms are also involved in YTHDC1-dependent SGK1 gene transcription still needs further exploration. More importantly, YTHDC1 is not the only m^6^A reader that mediates gene transcription. For instance, the m^6^A reader FXR1 recruits DNA 5mC dioxygenase to m^6^A-modified transcripts, increasing regional chromatin accessibility and in turn activating transcription ^36^. The m^6^A binder hnRNPG combines with m^6^A on RNA 5’ ends and protects nascent transcripts from integrator-mediated termination, thus enhancing transcription ^37^. Therefore, we cannot exclude other readers involved in SGK1 gene transcription. Consistent with these findings, our reporter gene assay revealed that the mutation of m^6^A motifs in the SGK1 transcript and deletion of the YTHDC1 binding region in the SGK1 gene promoter did not fully reverse METTL3 silencing-induced repression of reporter gene transcription. This finding suggests the potential presence of alternative regulatory pathways through which METTL3 influences SGK1 expression, possibly relying on other readers to promote transcription rather than relying solely on YTHDC1. Although the mechanism by which SGK1 gene transcription is regulated by mRNA m^6^A modification was mainly explored using rat SGK1 transcripts, our cross-species mRNA sequence alignment analysis revealed that the m^6^A motifs were relatively conserved in the mRNA sequences of mice and humans, suggesting the potential existence of similar regulatory mechanisms for SGK1 mRNA m^6^A modification and gene transcription across distinct species.

Currently, therapy aimed at inhibiting VSMC proliferation is a well-established approach for treating restenosis. The most prominent drugs used in drug-eluting stents thus far are paclitaxel (a microtubule stabilizing agent) and sirolimus (an mTOR inhibitor) ^1^. Our present study underscores the importance of METTL3-catalyzed m^6^A modification in promoting VSMC proliferation and contributing to in-stent restenosis. Encouragingly, of the recently developed small-molecule inhibitors targeting METTL3 for acute myeloid leukemia treatment ^38^ also possibly have potential for suppressing VSMC proliferation. Consequently, the introduction of stents eluted with METTL3 inhibitors may offer an innovative extension to current therapeutic strategies for managing restenosis following angioplasty and stenting.

## Nonstandard Abbreviations and Acronyms

AAV: adeno-associated virus
ALKBH5: alkB homolog 5 RNA demethylase
BrdU: 5-bromodeoxyuridine
CCK-8: cell counting kit-8
CABG: coronary artery bypass grafting
CEA: carotid endarterectomy
ChIP: chromatin immunoprecipitation
EdU: 5-Ethynyl-2’-deoxyuridine
FBS: fetal bovine serum
FLT1: fms related receptor tyrosine kinase 1
FTO: fat mass and obesity-associated protein
GAPDH: glyceraldehyde-3-phosphate dehydrogenase
GEO: gene expression omnibus data base
H&E: hematoxylin and eosin
IF: immunofluorescence
IRF1: interferon regulatory factor 1
m^6^A: N^6^-methyladenosine
MeRIP: m^6^A-specific methylated RNA immunoprecipitation
METTL3: methyltransferase-like 3
METTL14: methyltransferase-like 14
NOTCH3: neurogenic locus notch homolog protein 3
PCI: percutaneous coronary intervention
PDGF-BB: platelet derived growth factor BB
RT-qPCR: reverse transcription-quantitative real-time polymerase chain reaction
SGK1: serum/glucocorticoid regulated kinase 1
SMA: smooth muscle actin
TRAF2: TNF receptor-associated factor 2
TSS: transcriptional start site
VSMC: vascular smooth muscle cell
WTAP: Wilms tumor 1-associated protein
YTH: YT521-B homology
YTHDC1: YTH domain-containing protein 1

## Acknowledgments

We sincerely acknowledge Prof. Wengong Wang from Peking University for kindly providing the *Mettl3*^flox/flox^ mice. We thank Dr. Cihang Liu from Peking University for kindly providing technical support in the MeRIP-qPCR experiments. J.H. and Q.F. performed the experiments and analyzed the results. Z.D. and Z.Li. supported the transgenic mice experiments. Y.L., Z.Liu. and Y.Z. acquired and analyzed MeRIP sequencing and RNA sequencing. R.X. provided human samples. Q.D., X.Y. and F.Y. performed the immunofluorescence staining experiments. Y.J., W.K., H.T and Y.F. conceived the study, designed the experiments and analyzed the results. J.H., Q.F., H.T and Y.F. wrote the manuscript. Y.Z., Y.J., H.T., W.K. and Y.F. edited the manuscript. All authors have discussed the results and had the opportunity to comment on the manuscript.

